# Disease Severity Across Psychiatric Disorders Is Linked to Pro-Inflammatory Cytokines

**DOI:** 10.1101/2025.03.28.645923

**Authors:** Pierre Solomon, Monika Budde, Mojtaba Oraki Kohshour, Kristina Adorjan, Maria Heilbronner, Alba Navarro-Flores, Sergi Papiol, Daniela Reich-Erkelenz, Eva C. Schulte, Fanny Senner, Thomas Vogl, Lalit Kaurani, Dennis M. Krüger, Farahnaz Sananbenesi, Tonatiuh Pena, Susanne Burkhardt, Anna-Lena Schütz, Ion-George Anghelescu, Volker Arolt, Bernhardt T. Baune, Udo Dannlowski, Detlef E. Dietrich, Andreas J. Fallgatter, Christian Figge, Georg Juckel, Carsten Konrad, Fabian U. Lang, Jens Reimer, Eva Z. Reininghaus, Max Schmauß, Carsten Spitzer, Jens Wiltfang, Jörg Zimmermann, André Fischer, Peter Falkai, Thomas G. Schulze, Urs Heilbronner, Jeremie Poschmann

## Abstract

**Importance:** Numerous studies indicate that the traditional categorical classification of severe mental disorders (SMD), such as schizophrenia, bipolar disorders, and major depressive disorders, does not align with the underlying biology of those disorders as they frequently overlap in terms of symptoms and risk factors.

**Objective:** This study aimed to identify transdiagnostic patient clusters based on disease severity and explore the underlying biological mechanisms independently of the traditional categorical classification.

**Design:** We utilized data from 443 participants diagnosed with SMD of the PsyCourse Study, a longitudinal study with deep phenotyping across up to four visits. We performed longitudinal clustering to group patients based on symptom trajectories and cognitive performance. The resulting clusters were compared on cross-sectional variables, including independent measures of severity as well as polygenic risk scores, serum protein quantification, miRNA expression, and DNA methylation.

**Results:** We identified two distinct clusters of patients that exhibited marked differences in illness severity but did not differ significantly in age, sex, or diagnostic proportions. We found 19 serum proteins significantly dysregulated between the two clusters. Functional enrichment pointed to a convergence of immune system dysregulation and neurodevelopmental processes.

**Conclusion:** The observed differences in serum protein expression suggest that disease severity is associated with the convergence of immune system dysregulation and neurodevelopmental alterations, particularly involving pathways related to inflammation and brain plasticity. The identification of pro-inflammatory proteins among the differentially expressed markers underscores the potential role of systemic inflammation in the pathophysiology of SMD. These results highlight the importance of considering illness severity as a core dimension in psychiatric research and clinical practice and suggest that targeting immune-related mechanisms may offer promising new therapeutic avenues for patients with SMD.

**Key points:** 

**Question:** Can analyzing symptom trajectories and cognitive profiles across diagnostic categories reveal clinically relevant subgroups in severe mental disorders?

**Findings:** In this longitudinal study of 443 individuals with severe mental disorders, two distinct clusters emerged, differing significantly in illness severity, with the more severe group displaying elevated pro-inflammatory serum proteins, suggesting an association between disease severity and inflammation.

**Meaning:** These findings suggest that transdiagnostic clustering clarifies shared mechanisms, underscores the importance of inflammation in severe mental disorders, and highlights a promising avenue for novel therapeutic approaches.

## Introduction

Severe mental disorders (SMD) such as schizophrenia (SCZ), schizoaffective disorder (SCZA), bipolar disorder (BD), and major depressive disorder (MDD), are complex and multifaceted syndromes that have common symptoms and risk factors(1). Many characteristics such as cognitive dysfunction, functional impairment, and psychiatric symptoms are shared among SMD. There is evidence from behavioral genetic and genome-wide association studies (GWAS) that SMD are heritable, with a common phenomic(2,3) and genomic basis(4). Different nosological frameworks have been proposed based on account for these commonalities, with RDoC and HiTOP(5) emphasizing dimensionality rather than categorical classification. Also, the DSM-5 is now emphasizing dimensionality in addition to diagnostic categories. Moreover, multiple studies have associated SMD with a dysregulation of the immune system. For example, SMD are marked by an augmentation of pro-inflammatory proteins in the blood (6,7). Treatments of pro-inflammatory cytokines can induce SMD symptoms(8). The link between the immune system and SMD is so important that those disorders are recognized as neuroimmune disorders (9–11). However, a challenge that remains for clinicians, regardless of the diagnostic system used is how to differentiate individuals who will experience a severe form of psychopathology from milder cases. The gold standard are population-based longitudinal studies with a duration of several years or decades, ideally prior to disease onset. However, such studies are both extremely challenging and costly to conduct, and, in the end, most longitudinal studies have emphasized highly heterogeneous illness trajectories over time(12). Here, we take a different approach and instead focus on the short-term course of the five core aspects of SMD: we focus on these five core aspects, cognitive dysfunction, functional impairment, and psychotic, manic, and depressive symptoms, because they collectively represent the fundamental dimensions of impairment across diagnostic categories, allowing for a more nuanced and transdiagnostic perspective on the severity and course of SMD. We take advantage of The PsyCourse Study(13), a deeply phenotyped longitudinal sample of the psychotic-to-affective spectrum, collected specifically for this purpose. To identify clusters of patients that differ in their longitudinal course, we used *longmixr,* an R package that enables longitudinal clustering of mixed data types(14,15), specifically created for data collected in The PsyCourse Study. We identified two patient clusters that are similar in age, sex, and diagnostic proportions but exhibited distinct levels of severity. We then conducted comprehensive multi-omics analyses and found that proinflammatory cytokines are strongly associated with disease severity.

## Methods

### The PsyCourse Study

PsyCourse is a longitudinal, multisite, observational transdiagnostic study that was conducted in Germany and Austria(13). Participants provided written consent. Participants attended up to four visits evenly distributed over 18 months. With the aim to stratify participants independent of their diagnoses, we selected all participants diagnosed with SCZ, SCZA, BD, or MDD, according to DSM-IV, who participated in all four regular visits (443 participants).

### Longitudinal clustering

Longitudinal clustering was performed using the longmixr R package (Figure 1A)(14). First, longitudinally measured phenotypic data were meaningfully divided into five groups, based on prior knowledge, representing different domains of SMD, namely depressive, SCZ, manic, and cognitive symptoms, and global functioning.

a. Depressive symptoms (25 items): Rated on the Inventory of Depressive Symptomatology (IDS-C_30)_ (16). We excluded the items 11 to 14 as their values were missing for most participants.
b. Manic symptoms (11 items): Rated on the Young Mania Rating Scale (YMRS) (17)
c. SCZ symptoms (30 items): Rated on the Positive and Negative Syndrome Scale (PANSS), subscales positive symptoms, negative symptoms and general psychopathology (18)
d. Cognitive performance (5 items): Trail Making Test A and B (time), Verbal digit span forward, Verbal digit span backward, Digit-Symbol-Test
e. Functioning (6 items): Global Assessment of Functioning (GAF), along with the number of treatments prescribed and whether the participant was in a relationship

**Figure 1.**
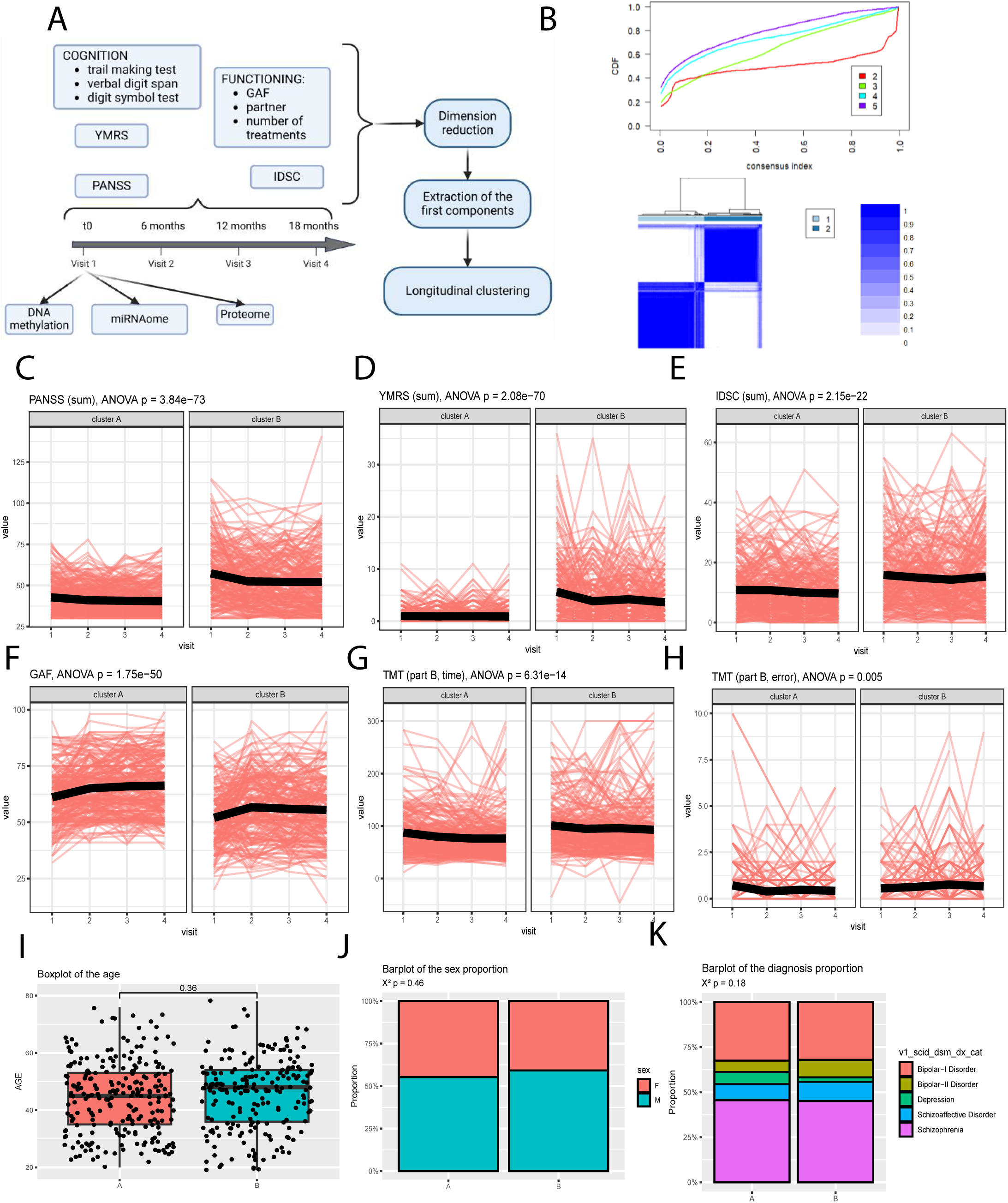
(A) Graph representing the strategy to classify the longmixr participants based on clinical evaluations. The consensus matrix displayed is the one computed by longmixr for the 2 clusters solution, while the consensus CDF was used to identify the optimal number of clusters. (B) Consensus CDF (upper) and consensus matrix (bottom) plots of our longmixr clustering. Spaghetti plots comparing the sum of the PANSS (negative, positive, and general) scales (C, ANOVA p-value < 0.001), the sum of the YMRS scale across the two clusters (D, ANOVA p-value < 0.001), the sum of the IDSC scale across the two clusters (E, ANOVA p-value < 0.001), the sum of the GAF scale across the two clusters (F, ANOVA p-value < 0.001), the time needed to finish Part B of the TMT tests across the two clusters (G, ANOVA, p-value < 0.001), and the number of errors made during the Part B of the TMT test (H, ANOVA p-value = 0.005). Boxplot comparing the age at the first visit of the two clusters (I, T Test p-value = 0.36), and barplots representing the sex (J, X² p-value = 0.458) and the diagnosis proportions in each cluster (K, X² p-value = 0.18).

Treatment-related variables (number of prescribed psychiatric treatments) were included within the functioning domain used for longitudinal clustering to partially account for treatment exposure.

Missing values were imputed when needed using multiple imputations (n=5 variables; R package *Amelia*). To reduce dimensionality within each group of variables, longmixr applies factorial analysis of mixed data (FAMD). The first component from each FAMD was corrected for the effects of age and sex, if necessary, and then used for longitudinal consensus clustering (Figure 1A). The longmixr clustering was performed with 1000 repetitions, and a seed value of 5114 algorithm for the final linkage. For the other settings, default values were used. The best solution was a two-cluster solution.

### PRS computation

Individuals were genotyped using the Illumina Infinium Global Screening Array-24 Kit (GSA Array). For details on QC see Solomon et al.). The PRS were computed using PRS-cs with default values (phi=auto settings) using the genotypes of the PsyCourse participants and GWAS summary of the trait studied that includes BD, MDD, SCZ, SCZ treatment resistance, and the P-Factor, a dimension of general psychopathology (19–21).

To assess whether genetics contributed to the differences in clusters, we compared PRS for MDD, BD, SCZ and for antipsychotic treatment resistance and the P-factor (19,22,23). Genotyping was described previously (24).

### Phenotypic characterization of the clusters

To validate the clustering, we compared the clusters for the sums of the scales and the scores of neuropsychological tests used for clustering at each visit. To ensure that there were no biases, we checked for differences in age, sex, and diagnosis.

Opcrit score: Operational Criteria Checklist for Psychotic and Affective Illness (OPCRIT) is a standardized diagnostic that provides detailed symptom ratings and operationally defined diagnoses based on established criteria (25). At the fourth visit OPCRIT item 90 was rated, assessing the disease course on an ordinal scale from “single episode with good remission” to “ongoing chronic disease with deterioration” of the psychiatric disorder diagnosed at the first visit of the PsyCourse Study. As this score was measured at a single visit, we didn’t use it to realize the clustering of the participants.

We compared the OPCRIT score and 2 other severity indicators that were not used for clustering, the body mass index (BMI), and childhood trauma using Childhood Trauma Screener (26), as well as 24 items, measured at the first visit, indicating the presence of somatic diseases. Statistics included X² tests for categorical variables, T-tests for single measurements, and ANOVA for multiple time-points. Analysis and figures were realized with R version 4.4.1.

### Somatic Disease comparison across the two clusters

We compared the presence of somatic disease (evaluated at the first visit of the PsyCourse Study) such as infectious diseases, Parkinson syndrome, allergies, … (See the Supplementary Table II) using X² test. The p-value of the tests were adjusted using the Benjamini-Hochberg method.

### Multiomics analysis

Propensity score matching was used to control for age, sex, and diagnosis when selecting participants across both clusters for the proteomic and methylome analysis. Serum from the first visit was used to quantify 384 proteins in 176 participants using Olink Explore Inflammation I panel. The selection of 176 participants was determined by the available capacity of two Olink Explore plates, each accommodating 88 samples plus 8 internal controls within a standard 96-well format, for a total of 176 samples across two plates. The *OlinkAnalyze* package was used for QC. For DNA methylation analysis 192 participants were analyzed (Illumina EPIC V2). The number of 192 participants was selected to match the proteomics sample size as closely as possible while fitting the array design of the EPIC V2 chip (8 samples per array). For the miRNAome analysis we used was sequenced using whole blood sample obtained on the first visit of the PsyCourse Study and sequenced in a previous study (27).

#### Proteomic data analysis

To identify differentially expressed proteins between the high-severity and low-severity clusters, we performed linear modeling using the limma package in R, including age as a covariate in the design matrix to adjust for its potential confounding effects on protein expression. Following model fitting, we applied the ashr package (adaptive shrinkage) to compute q-values and posterior effect sizes. This empirical Bayes approach stabilizes variance estimates and improves the estimation of significance and directionality in datasets with moderate sample sizes. As a quality control step, we excluded proteins with a posterior standard deviation (PosteriorSD) ≥ 0.02, to reduce the risk of overconfident shrinkage artifacts (28). Gene Ontology (GO) analysis was realized with the clusterProfiler R package on the 19 significant proteins to identify biological pathways.

Spearman correlations were computed between the normalized expression values of proteins (NPX) and variables from neuropsychological tests (Trail Making Test A (TMT-A) and B (TMT-B) (time), Verbal digit span forward (DGT_SP_FRW), Verbal digit span backward (DGT_SP_BW), Digit-Symbol-Test (DST)) and age of the participants. To account for age, we computed partial correlations using age as covariable. We adjusted the correlation p-values using the Benjamini-Hochberg method.

#### Methylome data analysis

The ages were imputed using the *methylclock* package and probes were filtered out if variance <0.05 resulting in 693 probes. Differential analysis was performed with *minfi* and *limma* packages while adjusting for age and sex.

#### miRNAome analysis

The miRNAome was obtained from the first visit and processed as previously described(24,27). *DESeq2* was used to compare the two clusters while adjusting for age, sex, and sequencing batches. Multiple hypothesis correction (Benjamini-Hochberg) was used.

#### Bioinformatics tools

Linkage Disequilibrium (LD) were computed using plink version 1.9. All analysis were performed with R (4.4.1). Packages used include:

- longmixr version 1.0.0
- DESeq2 version 1.44.0
- ggplot2 version 3.5.1
- OlinkAnalyze version 4.0.1
- FlowSorted.BloodExtended.EPIC version 1.1.2
- methylClock 1.10.0
- methylClockData 1.12.0
- minfi version 1.50.0
- limma version 3.60.6
- ashr version 2.2-63
- clusterProfiler version 4.12.6

## Results

### Longitudinal clustering of participants

Our aim was to find clusters of individuals independently of their ascertained DSM-IV diagnoses (Figure 1A, Methods, Table I). We explored different clustering solutions and used the cumulative distribution function (CDF) to determine the optimal number of clusters (Figure 1B). The CDF indicated that the two-cluster solution was optimal, as it demonstrated better binary separation characterized by a flatter curve with steep ascents at 0 and 1 (14).

**Table I:**
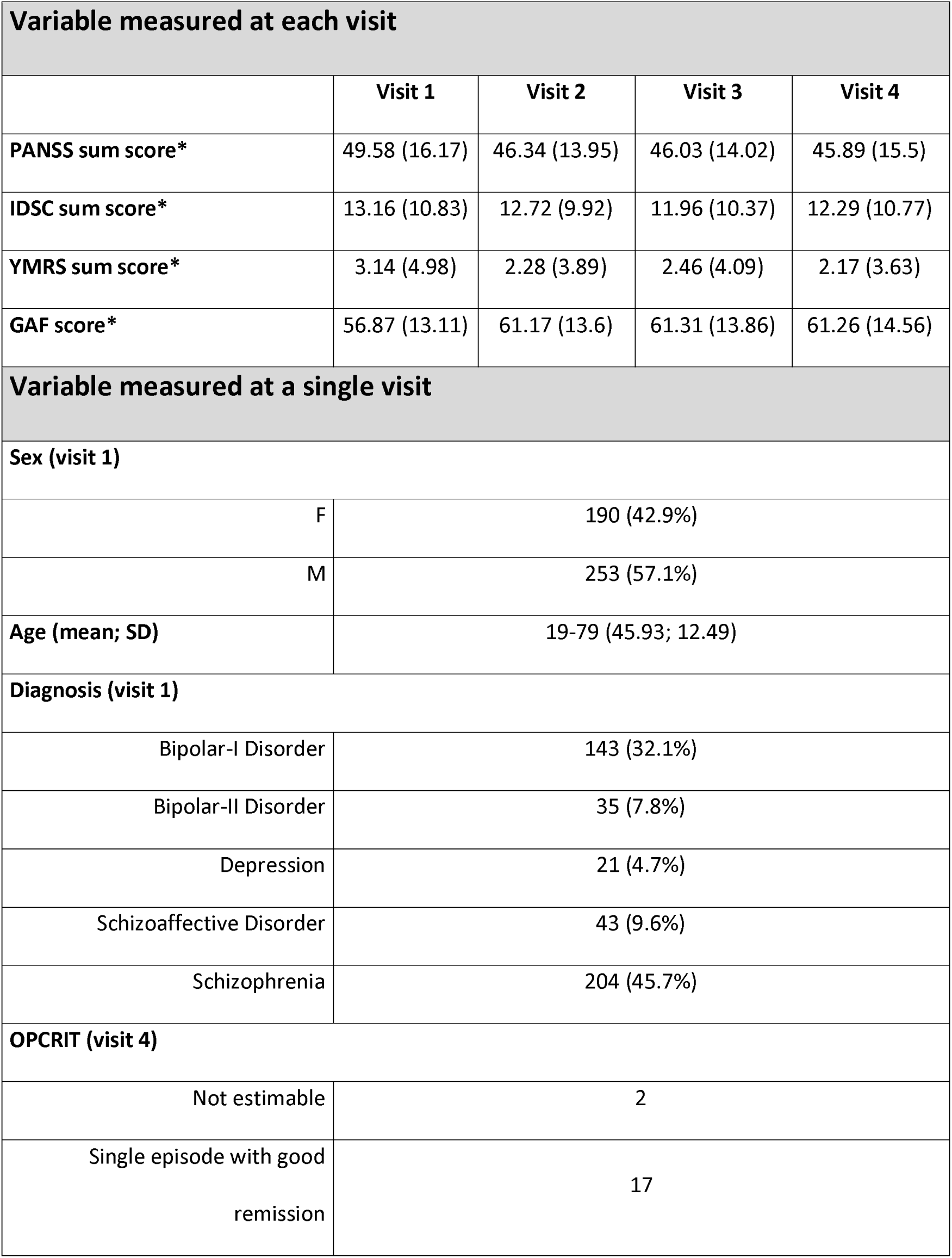
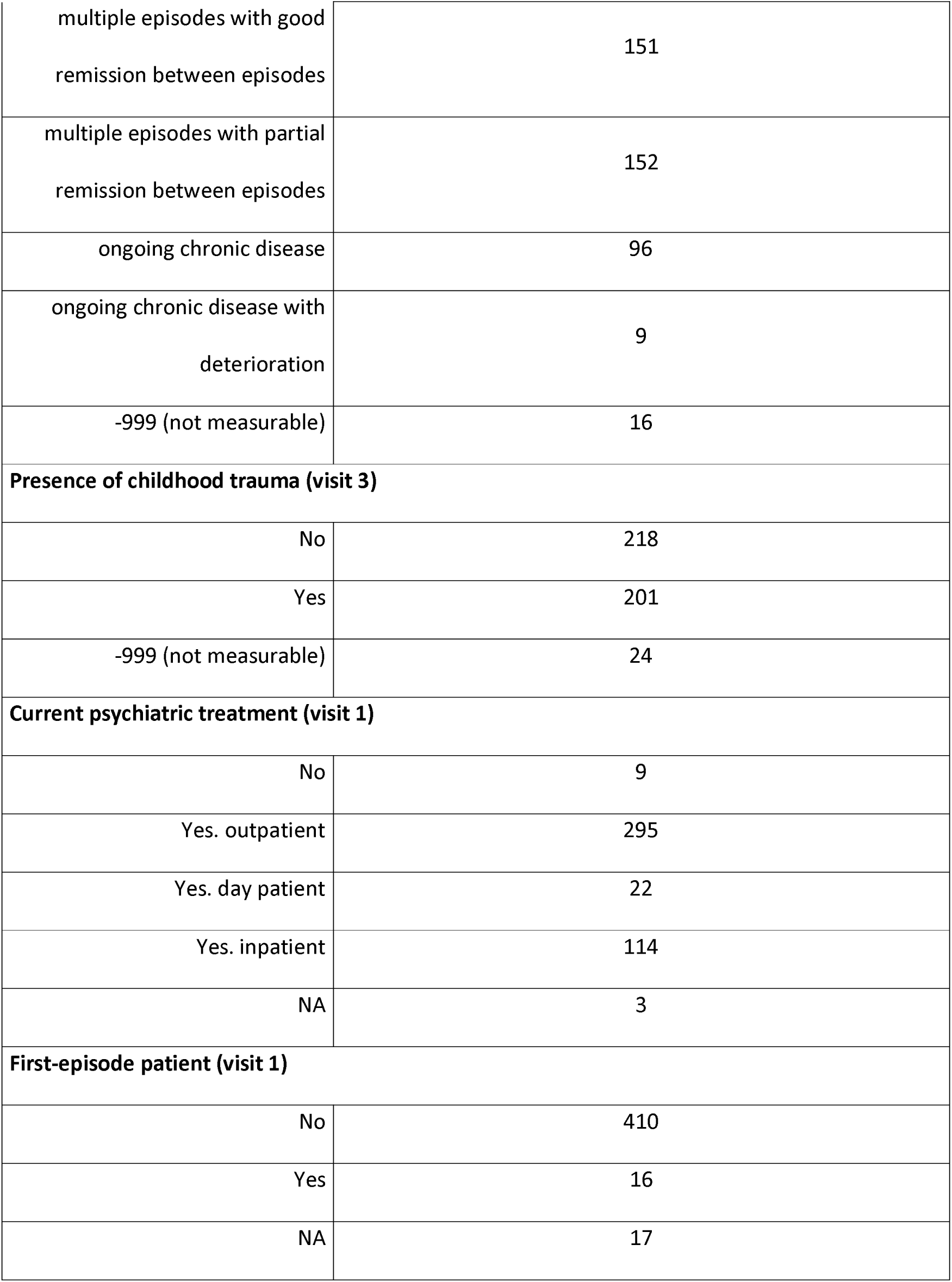
Variables measured by the PsyCourse Study. **Table that describe the participants included in our study with the sum of the medical scales measured at each visit of the PsyCourse Study including PANSS, IDSC, YMRS, and GAF scores. The table also indicate measures collected uniquely at the first visit (age, sex, diagnosis, if the disease just started, i.e. first-episode patients, and if the participant followed a psychiatric treatment). The presence of childhood trauma estimated at the third visit, and the disease course (OPCRIT) estimated at the fourth visit.**

| Variable measured at each visit |  |  |  |  |
| --- | --- | --- | --- | --- |
|  | Visit 1 | Visit 2 | Visit 3 | Visit 4 |
| PANSS sum score* | 49.58 (16.17) | 46.34 (13.95) | 46.03 (14.02) | 45.89 (15.5) |
| IDSC sum score* | 13.16 (10.83) | 12.72 (9.92) | 11.96 (10.37) | 12.29 (10.77) |
| YMRS sum score* | 3.14 (4.98) | 2.28 (3.89) | 2.46 (4.09) | 2.17 (3.63) |
| GAF score* | 56.87 (13.11) | 61.17 (13.6) | 61.31 (13.86) | 61.26 (14.56) |
| Variable measured at a single visit |  |  |  |  |
| Sex (visit 1) |  |  |  |  |
| F | 190 (42.9%) |  |  |  |
| M | 253 (57.1%) |  |  |  |
| Age (mean; SD) | 19-79 (45.93; 12.49) |  |  |  |
| Diagnosis (visit 1) |  |  |  |  |
| Bipolar-I Disorder | 143 (32.1%) |  |  |  |
| Bipolar-II Disorder | 35 (7.8%) |  |  |  |
| Depression | 21 (4.7%) |  |  |  |
| Schizoaffective Disorder | 43 (9.6%) |  |  |  |
| Schizophrenia | 204 (45.7%) |  |  |  |
| OPCRIT (visit 4) |  |  |  |  |
| Not estimable | 2 |  |  |  |
| Single episode with good remission | 17 |  |  |  |

|  |  |
| --- | --- |
| multiple episodes with good remission between episodes | 151 |
| multiple episodes with partial remission between episodes | 152 |
| ongoing chronic disease | 96 |
| ongoing chronic disease with deterioration | 9 |
| -999 (not measurable) | 16 |
| **Presence of childhood trauma (visit 3)** |  |
| No | 218 |
| Yes | 201 |
| -999 (not measurable) | 24 |
| **Current psychiatric treatment (visit 1)** |  |
| No | 9 |
| Yes. outpatient | 295 |
| Yes. day patient | 22 |
| Yes. inpatient | 114 |
| NA | 3 |
| **First-episode patient (visit 1)** |  |
| No | 410 |
| Yes | 16 |
| NA | 17 |

Both clusters showed significant differences across various symptom scales and neuropsychological assessments. Cluster B exhibited significantly higher SCZ symptom scores (PANSS, ANOVA **F**(1, 1768) = 359.59, p-value = 3.84e-73), manic symptoms (YMRS, ANOVA F(1, 1768) = 344,54, p-value = 2.08e-70), and depressive symptoms (IDS-C, ANOVA F(1, 1768) = 97,38, p-value = 2.15e-22) compared to Cluster A (Figure 1C–E). Compared to cluster A, cluster B also showed poorer functioning scores (GAF, ANOVA F(1, 1768) = 238, 05, p-value = 1.75e-50; Figure 1F).

In addition, patients in Cluster B performed worse for cognition, showing longer completion times on the Trail Making Test Part B (TMT-B, ANOVA F(1, 1768) = 57,19, p-value = 6.31e-14) and higher error rates (ANOVA F(1, 1615) = 7.898, p-value = 0.005), and had lower scores for Verbal Digit Span forward (ANOVA F(1, 1762) = 93,77, p = 4.85e-07) and backward (ANOVA F(1, 1768.22e-21) = 52.67, p = 5.87e-13), and Digit-Symbol Test (ANOVA F(1, 1762) = 93.77,p-value = 0.001; Figure 1G-H; Supplementary Figures 1A-E). In summary, Cluster B consists of participants with higher illness severity compared to cluster A.

Our clustering algorithm used longitudinal data from these symptom scales and neuropsychological tests to group patients based on their symptom trajectories and cognitive performance over time. Since the clusters were derived by analyzing patterns in these specific measures, we expected that the resulting clusters would differ significantly on these variables. However, it is noteworthy that the differences across these diverse scales and tests converge in the same direction, consistently indicating greater severity in Cluster B. The convergence of these independent measures across symptoms and cognition supports the conclusion that clustering differentiates patients primarily by severity.

### Comparison of Cross-Sectional Phenotypic Variables

Although age and sex were regressed out for clustering, this procedure does not constrain the resulting clusters to be balanced on these variables. We therefore examined age and sex distributions across clusters and found no significant differences in age (Figure 1I; mean age Cluster A: 44 years, Cluster B: 45 years; *t*-test *p*-value = 0.36) or sex (Figure 1J; Cluster A: 55,3% male, Cluster B: 40.8% male; χ² *p*-value = 0.46). Importantly, the clusters did not differ significantly in the proportions of diagnoses among SCZ, SCZA, BD, and MDD (Figure 1K; χ² *p*-value = 0.18). Additionally, the proportions of male and female subjects in the clusters were also similar for each individuals diagnosis (Supplementary Figure 1F and Supplementary Table I, X² adjusted p-value = 1 for each comparison). Age was also similar when stratified by sex and diagnosis individually, indicating that age and sex are distributed similarly in both clusters (Supplementary Figure 1G).

We next assessed whether somatic comorbidities differed between severity clusters using data collected at the first visit of the PsyCourse Study. Statistically significant differences were observed for diabetes, hypertension, and elevated cholesterol or triglyceride levels, but not for infectious diseases (Supplementary Figure 1F, Supplementary Table II, X² adjusted p-value < 0.05). These metabolic conditions are well-established comorbidities of SMD and are often linked to obesity and metabolic syndrome, both of which are exacerbated by antipsychotic treatment. Importantly, while the number of prescribed psychotropic medications was included as part of the clustering features, these somatic comorbidities emerged as independent markers of severity, consistent with previous reports linking cardiometabolic risk to worse clinical outcomes in psychiatric populations (29–33). Overall, cross-sectional variables such as age, sex, diagnosis, and treatment exposure were well balanced between clusters, supporting the robustness of the severity-based clustering approach.

### Comparison of Polygenic Profiles

To assess whether genetic factors contribute to the differences between our clusters, we used several PRS (see methods). We did not find significant differences between the clusters regarding PRS for MDD (Figure 2A; *t*-test adjusted *p*-value = 0.15), BD (Figure 2B; *t*-test adjusted *p*-value = 0.674), or SCZ (Figure 2C; *t*-test adjusted *p*-value = 0.894). Similarly, the sum of the PRS for these three disorders did not differ significantly between the clusters (Figure 2D; *t*-test adjusted *p*-value = 0.674). Furthermore, no significant differences were observed for SCZ treatment resistance PRS (Figure 2E; adjusted *p*-value = 0.674) or for the P factor PRS (Figure 2F; *t*-test adjusted *p*-value = 0.971). These results suggest that the observed differences between the clusters are not driven by common genetic risk factors captured by current PRS models.

**Figure 2.**
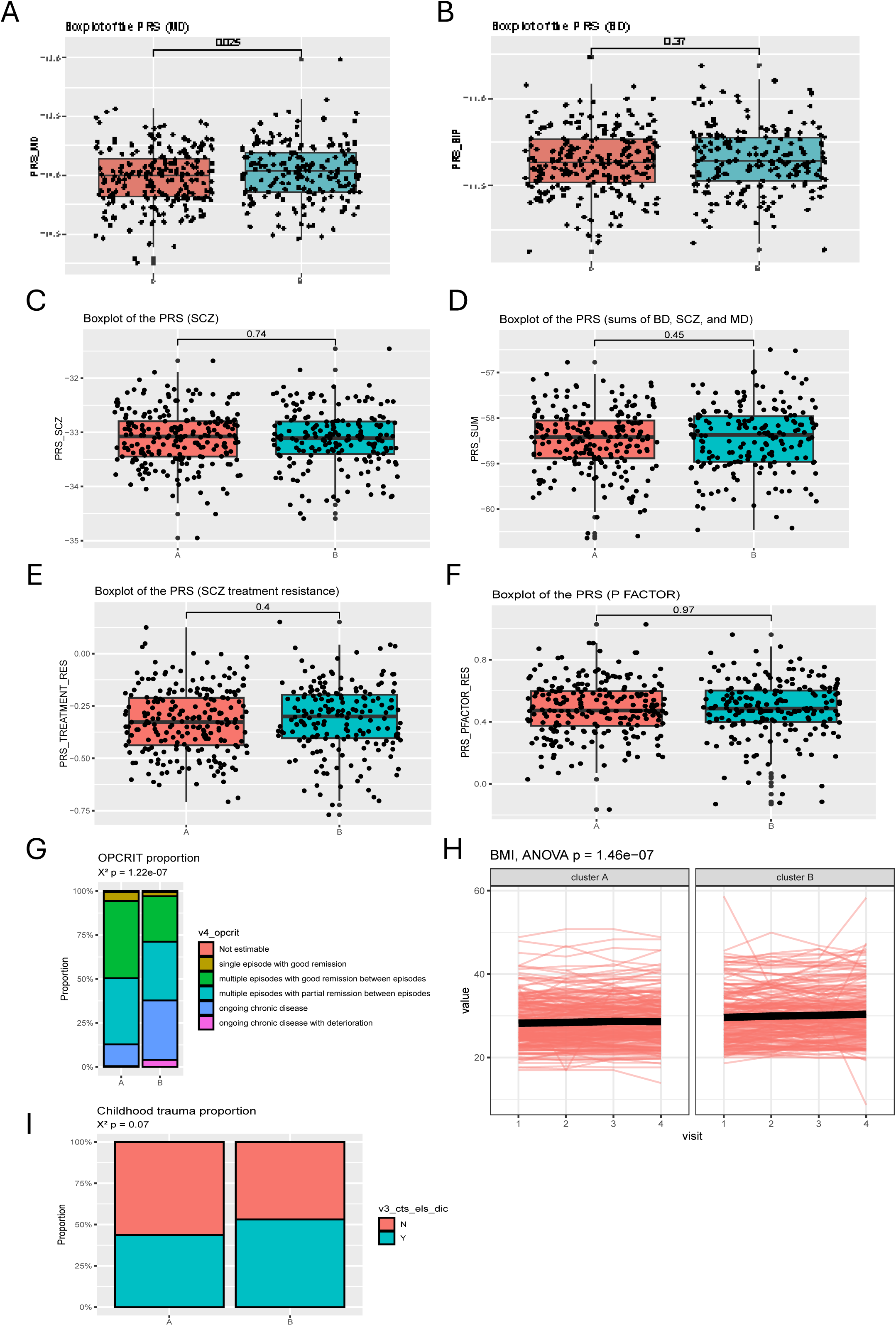
Boxplot comparing PRS between the two clusters, namely PRS for MD (A, T test adjusted p-value = 0.150), BD (B, T test adjusted p-value = 0.674), SCZ (C, T test adjusted p-value = 0.894), all 3 disorders (D, T test adjusted p-value = 0.674, sum of the individual PRS), SCZ treatment resistance (E, T test adjusted p-value = 0.674) and for the P factor (F, T test adjusted p-value = 0.971). Barplot representing the proportion of OPCRIT categories measured at the fourth visit of the PsyCourse study in each cluster (G, X² p-value < 0.001). Spaghetti plot comparing the BMI of the two clusters (H, ANOVA p-value < 0.001), and barplot representing the proportion of childhood trauma presence in each cluster (I, X², p-value = 0.07)

### Independent Measures of Severity

To further characterize the differences in severity between the clusters, we analyzed independent measures not used in the clustering process. We found a highly significant difference in the distribution of OPCRIT categories (χ² p-value= 1.22e-07; Figure 2G), with Cluster B showing a higher proportion of patients experiencing partial remission or ongoing chronic illness compared to Cluster A. This is an independent confirmation of a more severe illness trajectory in Cluster B.

BMI was also significantly higher in Cluster B compared to Cluster A (first visit mean BMI Cluster A: 28.22, first visit mean BMI Cluster_B: 29.6; ANOVA F(1, 1729) = 27.88, p-value = 1e-07; Figure 2H), supporting an association between severity and metabolic dysfunction. However, there was no significant difference in the rate of BMI change over time between the clusters (t-test estimate = 0.07; p-value = 0.33).Additionally, there was a trend toward a higher proportion of participants with a history of childhood trauma (CHT) in Cluster B (χ² p-value = 0.07; Figure 2I), suggesting a possible link between early adversity and greater illness severity, although this did not reach statistical significance. Together, these independent measures, especially the OPCRIT category distribution and BMI differences, consistently reinforce that Cluster B represents patients with a more severe and persistent illness course compared to Cluster A.

#### Multi-omics Analysis

To identify associated molecular alterations between the clusters we carried out 3 distinct molecular profiling approaches during the first visit for each patient group (Figure 3A).

**Figure 3.**
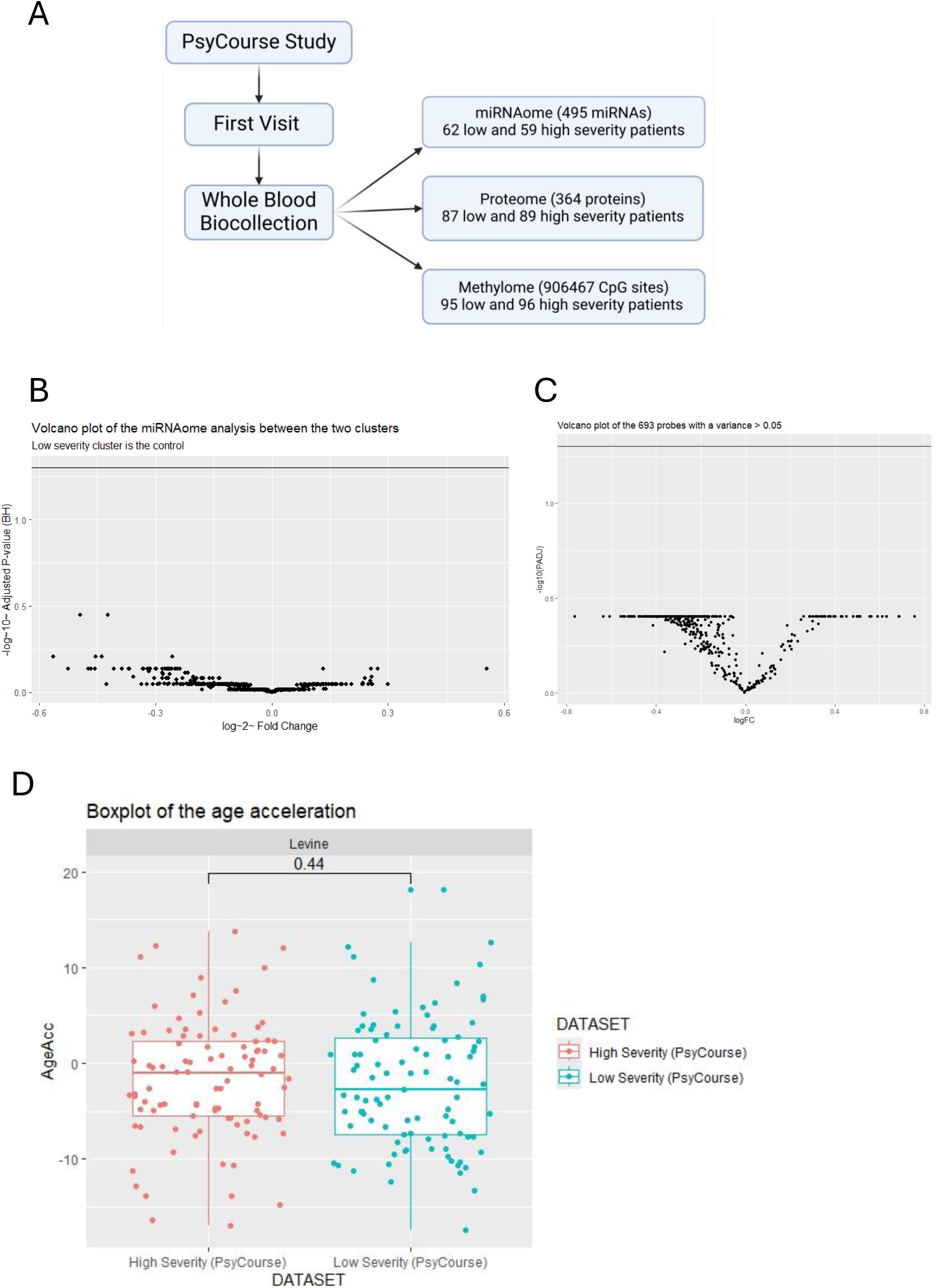
Flowchart that describes the multiomics analyses that were performed in this study (A) and volcano plot of the analysis comparing the miRNAome (B) and the methylome (C) of the two clusters. Boxplot comparing the age acceleration (imputed age – real age) of the two clusters (D).

#### miRNA Expression and Methylome Comparisons

We compared the miRNA profiles from the first visit of the PsyCourse Study of 125 participants (64 low-severity patients and 61 high-severity patients). PCA analysis did not reveal any major differences or batch effects (Supplemental figure 2A). Furthermore, none of the 465 miRNAs analyzed were differentially expressed between the two groups (adjusted p-values > 0.05 for all; Figure 3B), suggesting that miRNA expression profiles are unlikely to underlie the observed differences between the clusters. Similarly, we explored the methylomes of 192 patients using DNA samples (see supplemental figure 2B). We performed differential methylation analyses on the top variable methylation sites (see method). No CpG sites were significantly differentially methylated between the two clusters (adjusted p-value > 0.05 for all; Figure 3C). To further investigate a possible relationship between DNA methylation and severity we explored whether epigenetic age acceleration was distinct between the clusters. The imputed epigenetic age correlated strongly with the actual chronological age (Supplementary Figure 2C, Pearson’s R² = 0.83, p-value = 2.07e-75), but no significant difference was found between the clusters in epigenetic age acceleration (Wilcoxon test p-value = 0.44; Figure 3D).

#### Proteomic Differences in a targeted inflammatory protein Panel

We profiled 363 circulating serum proteins using the Olink Explore Inflammation panel to investigate molecular differences between severity clusters (Supplementary Figure 3A). Differential expression analysis was performed using the limma package, adjusting for age as a covariate (see methods). This analysis identified 19 differential proteins (adjusted p-value ≤ 0.05; Figure 4A, Supplementary Table III), indicating robust associations between protein expression and cluster membership. Although statistically significant, the differences in expression levels between clusters were modest in magnitude, as illustrated in the boxplots of the top four differentially expressed proteins (Figure 4B). This visual subtlety is expected given that the model accounts for inter-individual variation due to age, which reduces apparent differences while increasing statistical power.

**Figure 4.**
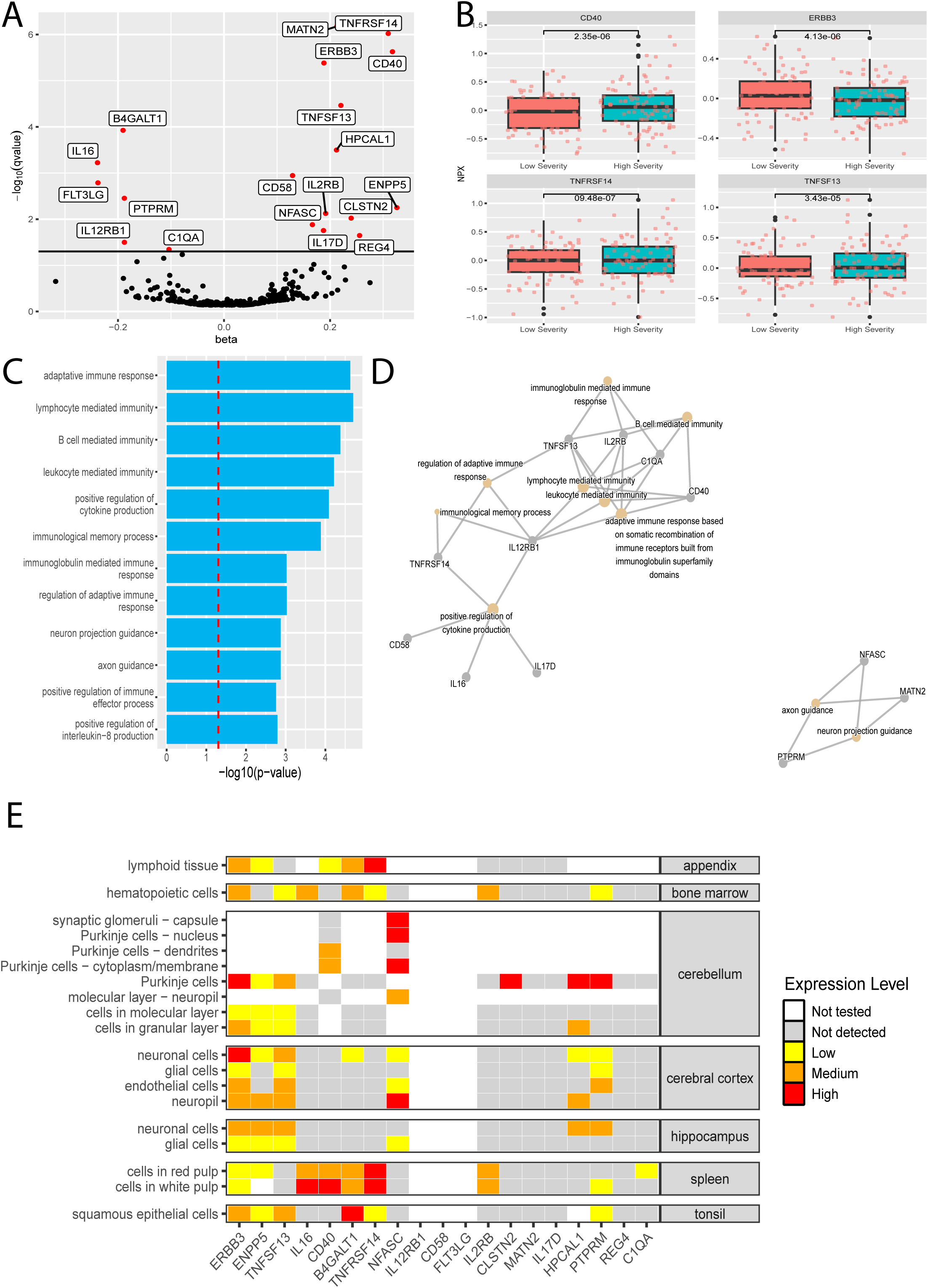
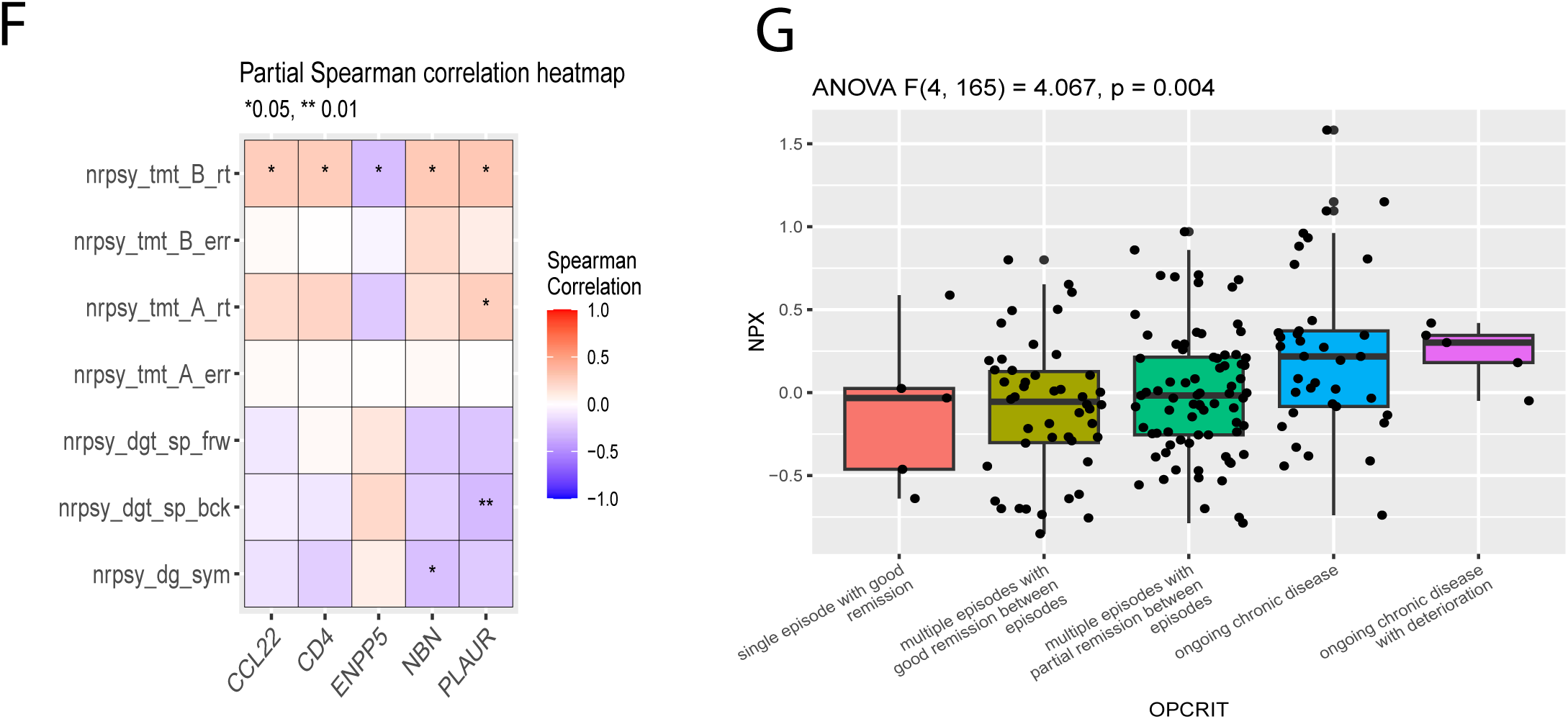
Volcano plot of the proteomicanalysis(A), and expression boxplot of the top 4 dysregulated proteins across the two clusters (B). Barplot of the p-values of the 12 significant GO terms (C) Network graph of the GO process enriched in the dysregulated proteins (D) in the high severity clusters. The color and size of the nodes correspond respectively to the p-value and the number of genes associated to the GO terms. The size of the edges represents the number of proteins in common between the GO terms. heatmap representing the expression levels of the dysregulated proteins in brain and immune cell-types according to the Human Protein Atlas (E). Correlation heatmap of the partial correlation of spearman between the protein’s expression and the neuropsychological assessment variables (* p<0.05, ** p<0.005). Boxplot representing the expression of the PLAUR gene according to the different OPCRIT categories (G

To ensure that pharmacological treatments did not confound the observed differences in protein expression, we assessed the relationship between protein levels and treatment exposure at baseline. Specifically, we computed Spearman correlations between the NPX values of the Olink panel proteins, and the number of psychotropic medications prescribed at the first visit (including antipsychotics, antidepressants, mood stabilizers, and tranquilizers). After adjusting p-values for multiple comparisons using the Benjamini-Hochberg method, no significant correlations were identified (Supplementary Table IV, Supplementary Figure 4), suggesting that treatment burden did not account for the proteomic differences observed between severity clusters.

A literature review revealed that 9 of the 19 differentially expressed proteins (47.3%) have been previously associated with psychiatric disorders or related neurological processes such as neuroplasticity, compared to 82 of the remaining 344 proteins (23.8%) (Supplementary Table V). This represents a significant enrichment of psychiatric-related proteins among the dysregulated set (Fisher’s exact test, p-value = 0.0293).

Gene Ontology (GO) enrichment analysis of differentially expressed proteins revealed that the high-severity cluster exhibited significant enrichment in immune-related pathways (adjusted p-value ≤ 0.05; Figure 4C and 4D, Supplementary Table VI). The top enriched processes included “adaptive immune response,” “lymphocyte mediated immunity,” “B cell mediated immunity,” and “positive regulation of cytokine production,” all of which suggest an overactive immune signaling landscape. These pathways were connected in a highly coherent network structure, indicating shared upstream regulation and overlapping molecular effectors. Key hub proteins associated with this immune activation included IL2RB, CD40, C1QA, and TNFRSF13, each of which has been previously implicated in immune cell signaling and neuroimmune interactions. Beyond the immune component, GO terms related to neural development and organization were also enriched in the high-severity group. These included “axon guidance” and “neuron projection guidance,” suggesting that proteins involved in CNS structural plasticity and connectivity are also dysregulated. The gene products implicated in this module (e.g., NFASC, PTPRM, MATN2) play key roles in axon pathfinding and myelination, processes that are critical for maintaining functional brain circuits. These findings point toward a dual signature of severity: an elevated inflammatory profile involving adaptive immunity, and alterations in neurodevelopmental pathways that may affect cognitive and emotional regulation. Importantly, these changes appear in serum, underscoring the relevance of peripheral biomarkers for capturing central pathophysiology. Interestingly, these two functional categories were driven by distinct subsets of proteins, suggesting that both immune and neurodevelopmental pathways may independently contribute to the biological differences between severity clusters.

To better understand the tissue origin of these differentially expressed proteins, we examined their expression patterns using the HumanProtein Atlas database (34) (Figure 4E). This analysis confirmed the neuro-immune dichotomy: many of the proteins upregulated in high-severity patients were either of hematopoietic origin, with high expression in lymphoid tissues such as the spleen, tonsils, and bone marrow, or of neuronal origin, with specific expression in the cerebellum, cerebral cortex, and hippocampus. For instance, proteins such as TNFRSF13, IL2RB, and CD40 were enriched in immune tissues, while NFASC, PTPRM, and MATN2 were predominantly expressed in neural compartments. This spatial expression pattern supports a neuroimmune interface model in which disease severity is linked to cross-talk between the immune system and the brain. The simultaneous enrichment suggests that systemic inflammation may be the cause or consequence of neuronal integrity in high-severity psychiatric patients.

To extend on this and explore the potential link between protein expression and cognitive function, we performed Spearman and partial Spearman correlation analyses between the expression levels of all expressed proteins and neurocognitive assessment scores from the first visit of the PsyCourse study (Supplementary Figure 3B; Figure 4F). The partial correlations were adjusted for age to identify associations independent of this confounder. This analysis identified five proteins (CCL22, CD4, ENPP5, NBN, and PLAUR) that showed significant age-independent correlations with at least one cognitive variable (adjusted p<0.05). All five proteins were associated with performance on the Trail Making Test Part B (TMT-B, nrpsy_tmt_B_rt): ENPP5 was negatively correlated (indicating better performance with higher expression), while CCL22, CD4, NBN, and PLAUR were positively correlated (indicating worse performance with higher expression).

In addition, NBN was negatively correlated with Digit-Symbol Test scores (nrpsy_dg_sym), and PLAUR showed a negative correlation with Verbal Digit Span forward (nrpsy_dgt_sp_frw) and a positive correlation with Trail Making Test Part A time (nrpsy_tmt_A_rt). These findings suggest that ENPP5 expression is positively associated with cognitive performance, whereas higher expression of CCL22, CD4, NBN, and PLAUR is associated with poorer performance across several cognitive domains. PLAUR showed the strongest and most consistent correlations and has previously been linked to cognitive outcomes in independent studies, supporting its relevance as a candidate marker of cognitive dysfunction. Interestingly, PLAUR expression levels varied significantly across clinical disease course categories as defined by OPCRIT (ANOVA F(4,165) = 4.067, p = 0.004), with expression progressively increasing from single-episode remission to ongoing chronic disease with deterioration (Figure 4F). This pattern suggests that higher PLAUR expression is associated with more severe and persistent forms of psychiatric illness.

## Discussion

Our study leverages comprehensive longitudinal phenotyping over four visits to demonstrate that disease severity in SMD can be characterized independently of traditional diagnostic categories. By employing a transdiagnostic clustering approach, we identified transdiagnostic patient groups that differ primarily in illness severity. This approach allowed us to study the symptoms across disorders without relying on traditional categorical diagnoses. While there were no significant differences in diagnosis, age and sex overall, we did observe a significant difference in the proportion of female patients with SCZA disorder between the clusters. This specific variance may be due to unique clinical features within this subgroup, but overall, the similar diagnostic proportions highlight that the clusters represent groups differentiated by severity across diagnostic categories.

This finding underscores the significant overlap in symptoms across disorders such as SCZ, SCZA, BD, and MDD, highlighting the limitations of categorical diagnoses in capturing the complexity of psychiatric conditions. The overlapping symptoms between different psychiatric disorders have been well-documented and suggest that common underlying mechanisms may contribute to disease manifestation(3,4). Cognitive dysfunction, functional impairment, and mood disturbances are a hallmark multiple diagnoses. Our results demonstrate that these shared symptoms coalesce to define severity levels that transcend diagnostic boundaries. This supports dimensional models of psychopathology, such as RDoC and HiTOP, which advocate for assessing mental disorders along continuous dimensions rather than discrete categories(35,36).

Importantly, we found that independent measures not used in the clustering process, especially the OPCRIT item and BMI also differed between the clusters, further confirming the robustness of our severity classification. The high-severity cluster had a higher proportion of patients experiencing partial remission or ongoing psychiatric chronic illness according to the OPCRIT item, indicating a more severe overall illness trajectory. Additionally, patients in the high-severity cluster exhibited higher BMI, which has been associated with increased psychiatric symptom severity, poorer cognitive performance and poorer clinical outcomes in patients with affective disorders (37,38). Weight gain is also a common side effect of antipsychotic treatment and is associated with poorer medication adherence in patients with SCZ, which increases the risk for recurrent illness episodes and poorer outcome long term(39). Although the difference in CHT was not statistically significant, there was a trend toward a higher proportion in the high-severity cluster, aligning with literature that associates CHT with more severe psychiatric outcomes(40,41). These independent measures, which were not part of the clustering variables, reinforce our finding that the clusters are differentiated primarily by severity.

The genetic analysis revealed no significant differences in PRS for SCZ, BP, or MDD between the high-severity and low-severity clusters. This suggests that genetic variants identified in GWAS may be more closely associated with disease susceptibility rather than the severity of illness. In psychiatry, the largest GWAS focus on the risk of developing a disorder by comparing cases to controls, which may not capture genetic factors influencing the course or severity of the disease once it manifests. It is also possible that rare variants, structural variations, or gene– environment interactions play a more significant role in determining illness severity.

Our proteomic analysis revealed that disease severity is associated with robust but modest proteomic changes spanning immune activation and neural development processes. Specifically, we identified two distinct GO modules, one related to the immune activation and another linked to axon guidance, a process essential for brain plasticity and synaptic connectivity and previously associated with psychiatric disease risk (42,43). This dual enrichment suggests that dysregulation of immunity and disrupted neurodevelopmental signaling may contribute to increased illness severity and altered brain function in SMD. Moreover, several proteins within the immune-related GO terms have been previously implicated in psychiatric pathogenesis, including IL2RB, CD40, C1QA, and members of the TNF superfamily (44–48). In parallel, neurodevelopmental proteins such as NFASC and PTPRM, enriched in the axon guidance GO term, are both associated with the genetic risk of SCZ and BD (49–51), reinforcing the neurobiological relevance of this signature.

Given that cognitive dysfunction is a critical aspect of disease severity, we investigated whether the expression levels in our panel were associated with cognitive performances. Using partial Spearman correlations adjusted for age, we identified five proteins (CCL22, CD4, ENPP5, NBN, and PLAUR) that showed significant associations with at least one cognitive variable (adjusted p < 0.05). It is noteworthy that some of these proteins have been previously associated with psychosis or nervous system development. For example, among these, ENPP5 was positively associated with cognitive performance, particularly on the Trail Making Test Part B (TMT-B), suggesting a potential protective role, in line with prior Mendelian randomization studies linking ENPP5 expression to intelligence (52). Conversely, higher expression of CCL22, CD4, NBN, and PLAUR was associated with poorer performance across several cognitive domains, including attention, executive function, and working memory (53,54). Among these, PLAUR emerged as the most consistently associated protein, showing multiple significant correlations with cognitive performance even after adjusting for age. PLAUR is expressed in microglia within the CNS and plays a role in synaptic remodeling and GABA-A receptor regulation(55). Moreover, the soluble form of PLAUR has been implicated in several neurological disorders, potentially serving as a marker of neuroinflammation and blood-brain barrier dysfunction(56,57). Importantly, PLAUR expression also tracked closely with clinical disease trajectory. When stratified by OPCRIT disease course categories, PLAUR levels showed a significant increase from individuals with single-episode remission to those with chronic and deteriorating disease. This progressive pattern links PLAUR not only to cognitive deficits but also to long-term illness severity, making it a promising transdiagnostic biomarker candidate. Collectively, these findings suggest that disease severity in SMD is not solely driven by brain-specific pathology but is instead shaped by the interplay between immune dysregulation and CNS connectivity. This neuroimmune interface may represent a tractable target for novel therapeutic interventions and severity biomarkers in psychiatry.

In contrast to the proteomic findings, our analysis revealed no significant differences in miRNA expression or DNA methylation profiles between the high-severity and low-severity clusters. miRNA expression and DNA methylation are regulatory mechanisms that may require longer timescales to manifest significant changes. These epigenetic modifications are often more stable and may not fluctuate significantly over the short-term course captured in our study(58). An important consideration is that our analysis of miRNA and DNA methylation were conducted using blood which may not capture alterations occurring in the brain. The brain and blood can have distinct epigenetic landscapes, and peripheral markers may not fully represent central nervous system processes.

### Limitations

Despite the strengths of our comprehensive multi-omics approach and longitudinal design, our study has several limitations. First, the use of peripheral blood samples for miRNA and DNA methylation analyses may not accurately reflect brain-specific epigenetic changes associated with psychiatric disorders or be a consequence of lack of statistical power due to the sample size. Second, due to a reasonable but limited sample size in a homogenous population it is relevant to replicate our findings in larger and heterogenous populations in future studies. Third, our study is based on the PsyCourse Study with a predominantly European ancestry. A future study that repeats our workflow should use cohort with a more diverse larger genetic ancestry. Fourth, our sample size may have been too limited in both of those analysis considering the substantial statistical power required to detect small effect sizes due to the high dimensionality of the data and the need for stringent multiple testing correction. Larger studies will be necessary to confirm that disease severity is not associated with changes in miRNA expression or DNA methylation. Fifth, although treatment-related variables were partially considered during clustering (e.g., number of prescribed treatments) and a PRS for treatment resistance was included in the analysis, specific treatment types, dosages, and durations were not systematically adjusted for the multi-omics comparisons. However, we conducted correlation analyses between the number of prescribed psychotropic medications at baseline and protein expression levels and found no significant associations after multiple testing correction. This suggests that treatment burden at the time of sampling is unlikely to explain the observed proteomic differences between severity clusters. Nevertheless, residual confounding due to unmeasured treatment variables, such as medication class effects, cumulative exposure, or treatment response, cannot be fully excluded. Future analyses incorporating detailed longitudinal treatment histories will be necessary to further address this limitation.

Sixth, a further limitation is the imbalance in diagnostic representation within the study cohort, with SCZ being the most frequent diagnosis. While our clustering approach did not incorporate diagnostic labels and clusters did not differ in diagnostic composition, we acknowledge that the sample composition may still introduce subtle biases in transdiagnostic interpretations and generalizability.

Seven, given the exploratory multiomics design, we opted to profile several molecular layers in parallel (genotypes, DNA methylation, miRNAome, and proteome), which necessarily limited the number of participants per assay. Future studies building on these findings may consider focusing on a single modality, such as proteomics, which showed the most robust associations in our study, to enable larger sample sizes and greater statistical power for detecting biologically meaningful effects.

### Clinical Relevance

Our findings have direct implications for how SMD might be better stratified and treated in clinical practice. By identifying transdiagnostic clusters based on symptom trajectories and cognitive performance—rather than diagnostic labels—we highlight the potential of severity-based classification for guiding personalized treatment approaches, in line with dimensional models of psychiatry such as RDoC and HiTOP frameworks (35,36).

The high-severity cluster exhibited a distinct pro-inflammatory proteomic profile, consistent with evidence linking systemic inflammation to worse psychiatric outcomes. Elevated inflammatory markers have been associated with treatment resistance, greater symptom severity, and cognitive impairment in patients with SCZ, BD, and MDD (59–61).

These findings support a growing body of research indicating that anti-inflammatory or immune-modulating treatments may offer clinical benefits, particularly as adjunctive therapies in patients with elevated inflammation. Meta-analyses and clinical trials have shown that Nonsteroidal anti-inflammatory drugs (e.g., celecoxib) and cytokine inhibitors may reduce depressive and psychotic symptoms in selected subgroups (62–64). In this context, evaluating inflammatory markers in routine psychiatric care could help stratify patients based on underlying biological risk and guide treatment intensity or augmentation strategies. Such approaches are increasingly being advocated as part of precision psychiatry models (65).

In conclusion, our study emphasizes the importance of comprehensive, longitudinal phenotyping in capturing the complexity of SMD beyond categorical diagnoses. Our multi-omics analysis further reveals that increased disease severity is associated with a pro-inflammatory proteomic profile independently of the clinical diagnoses, and potentially contributing to cognitive dysfunction and the severity of the psychiatric symptoms. Thus, our study highlights the potential of the immune proteome as candidate biomarkers of psychosis severity.

Our work underscores the importance of considering illness severity as a key dimension in psychiatric research and clinical practice. Tailoring interventions based on severity rather than diagnosis alone may improve patient outcomes, and targeting inflammatory pathways could offer new therapeutic avenues for patients with SMD.

## Supporting information

Supplementary Table III

## Acknowledgments

This work is part of the MulioBio project funded by a NEURON-ERANET grant (JTC2019) and Pierre Solomon thesis co-funded by the region Pays de la Loire, France (order n°2020_10085). We are most grateful to the Genomics Core Facility GenoA and France Genomique, member of Biogenouest and Institut Français de Bioinformatique (IFB) (ANR-11-INBS-0013) for the use of their resources and their technical support.

TGS and PF are supported by the Deutsche Forschungsgemeinschaft (DFG) within the framework of the projects www.kfo241.de and www.PsyCourse.de (SCHU 1603/4-1, 5-1, 7-1; FA241/16-1). TGS is further supported by the Dr. Lisa Oehler Foundation (Kassel, Germany), IntegraMent (01ZX1614K), BipoLife (01EE1404H), e:Med Program (01ZX1614K), GEPI-BIOPSY (01EW2005), MulioBio (01EW2009). This work was endorsed by German Center for Mental Health (DZPG; to PF and TGS, FKZ: 01EE2303A, 01EE2303F). UH was supported by European Union’s Horizon 2020 Research and Innovation Program (PSY-PGx, grant agreement No 945151) and the Deutsche Forschungsgemeinschaft (DFG, German Research Foundation, project number 514201724). EXC 2067/1 390729940. FS was supported by the GoBIO project miRassay from the BMBF. The study was endorsed by the Federal Ministry of Education and Research (Bundesministerium für Bildung und Forschung [BMBF]) within the initial phase of the German Center for Mental Health (DZPG) (grant: 01EE2303A, 01EE2303F to PF).

Kristina Adorjan, Monika Budde, Peter Falkai, Maria Heilbronner, Urs Heilbronner, Alba Navarro-Flores, Mojtaba Oraki Kohshour, Sergi Papiol, Daniela Reich-Erkelenz, Eva C. Schulte, Thomas G. Schulze, Fanny Senner, and Thomas Vogl are part of the PsyCourse core team.

Ion-George Anghelescu, Volker Arolt, Bernhardt T. Baune, Udo Dannlowski, Detlef E. Dietrich, Andreas J. Fallgatter, Christian Figge, Fabian Lang, Georg Juckel, Carsten Konrad, Jens Reimer, Eva Z. Reininghaus, Max Schmauß, Carsten Spitzer, Jens Wiltfang, and Jörg Zimmermann represent clinical centers involved in the recruitment and sample collection of the PsyCourse Study.

We would like to express our profound gratitude to all study participants without whom this work would not have been possible.

## Author contribution

Pierre Solomon: Validation, Formal Analysis, Investigation, Writing - original draft, Writing - review & editing, Visualization Monika Budde: Resources, Data curation, Writing - original draft, Writing - review & editing Mojtaba Oraki Kohshour: Resources, Data curation, Writing - review & editing Kristina Adorjan: Resources, Data curation, Writing - review & editing Maria Heilbronner: Resources, Data curation, Writing - review & editing Alba Navarro-Flores: Resources, Data curation, Writing - review & editing Sergi Papiol: Resources, Data curation, Writing - review & editing Daniela Reich-Erkelenz: Resources, Data curation, Writing - review & editing Eva C. Schulte: Resources, Data curation, Writing - review & editing Fanny Senner: Investigation, Resources, Data curation, Writing - review & editing Thomas Vogl: Resources, Data curation, Writing - review & editing Lalit Kaurani: Resources, Data curation, Writing - review & editing Dennis M. Krüger: Methodology, Resources, Data curation, Writing - review & editing Farahnaz Sananbenesi: Resources, Data curation, Writing - review & editing Tonatiuh Pena: Methodology, Resources, Data curation, Writing - review & editing Susanne Burkhardt: Resources, Data curation, Writing - review & editing Anna-Lena Schütz: Resources, Data curation, Writing - review & editing Ion-George Anghelescu: Resources, Data curation, Writing - review & editing Volker Arolt: Resources, Data curation, Writing - review & editing Bernhardt T. Baune: Resources, Data curation, Writing - review & editing Udo Dannlowski: Resources, Data curation, Writing - review & editing Detlef E. Dietrich: Resources, Data curation, Writing - review & editing Andreas J. Fallgatter: Resources, Data curation, Writing - review & editing Christian Figge: Resources, Data curation, Writing - review & editing Georg Juckel: Resources, Data curation, Writing - review & editing Carsten Konrad: Resources, Data curation, Writing - review & editing Fabian U. Lang: Resources, Data curation, Writing - review & editing Jens Reimer: Resources, Data curation, Writing - review & editing Eva Z. Reininghaus: Resources, Data curation, Writing - review & editing Max Schmauß: Resources, Data curation, Writing - review & editing Carsten Spitzer: Resources, Data curation, Writing - review & editing Jens Wiltfang: Resources, Data curation, Writing - review & editing Jörg Zimmermann: Resources, Data curation, Writing - review & editing André Fischer: Investigation, Resources, Writing - review & editing Peter Falkai: Resources, Writing - review & editing Thomas G. Schulze: Conceptualization, Investigation, Resources, Writing - review & editing, Funding acquisition Urs Heilbronner: Conceptualization, Investigation, Resources, Data curation, Writing - original draft, Writing - review & editing, Project administration Jeremie Poschmann: Conceptualization, Investigation, Resources, Data curation, Writing - original draft, Writing - review & editing, Supervision, Project administration, Funding acquisition

---

**Supplementary Figure 1**

Spaghetti plots showing the time needed to finish (A, ANOVA p-value < 0.001) and the number of errors (B, ANOVA p-value =0.146) made during Part A of the TMT test, as well as the scores of the Digit Symbol test (C, ANOVA p-value < 0.001), Verbal Digit Span test forward (D, ANOVA p-value < 0.001) and backward (E, ANOVA p-value < 0.001) in both clusters. (F) Barplots indicate the sex proportion in the two clusters across the different categorical diagnoses, and (G) Comparison of age between the clusters grouped by age and diagnosis using T tests (pvalues adjusted with BH method).

**Supplementary Figure 2**

PCA plot of the miRNAome expression data (A) of the methylation state of CpG site (A) of the patients. Correlation between real age of the participants during the first visit of the PsyCourse study and the DNA imputed age using the Levine model of the methylClock R package (C),

**Supplementary Figure 3**

PCA plot of the protein expression in NPX (A). Correlation heatmap of the partial correlation of spearman between the proteins’ expression of the entire panel and the neuropsychological assessment variables (B, * p<0.05, ** ^p<0.01, *** p<0.005, **** p<0.001). Heatmap representing the expression level of the dysregulated protein in all cell types available in the Human Protein Atlas (C).

**Supplementary Figure 4**

Correlation plot of the 19 significant proteins identified in our dysregulation analysis and the number of treatments prescribed during the first visit of the PsyCourse Study (the text in each scatter plot indicate the correlation and the non-adjusted p-values).

**Supplementary Table I**

Sex Proportions across the clusters for each diagnosis and results of their comparisons (X²).

**Supplementary Table II**

**Comparison of the proportions of patients with somatic disorders between the two clusters, assessed using** χ**² tests.Supplementary Table III**

Results of the limma analysis comparing the inflammatory proteome across the two clusters.

**Supplementary Table IV**

Results of the spearman correlation between the proteins’ NPX and the number of prescribed treatments at the first visit of the PsyCourse Study.

**Supplementary Table V**

Summary of whether the dysregulated proteins identified in our analysis have been previously associated with SCZ, BD, or MDD in our bibliographic review.

**Supplementary Table VI**

Results of the enrichment analysis using the dysregulated proteins

