## Supplementary figures and images for "Disease Severity Across Psychiatric Disorders Is Linked to Pro-Inflammatory Cytokines"

### Supplementary Figure 1

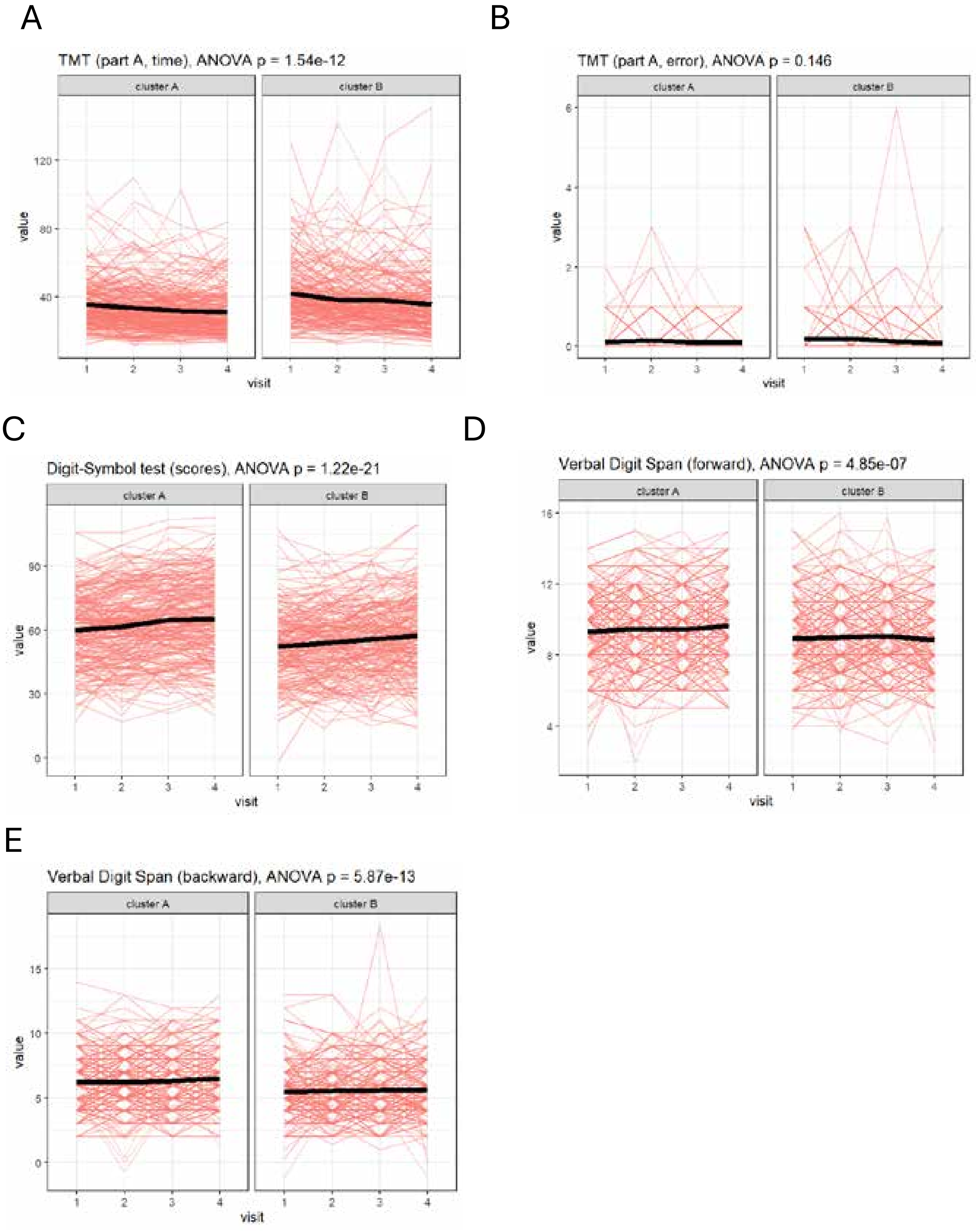

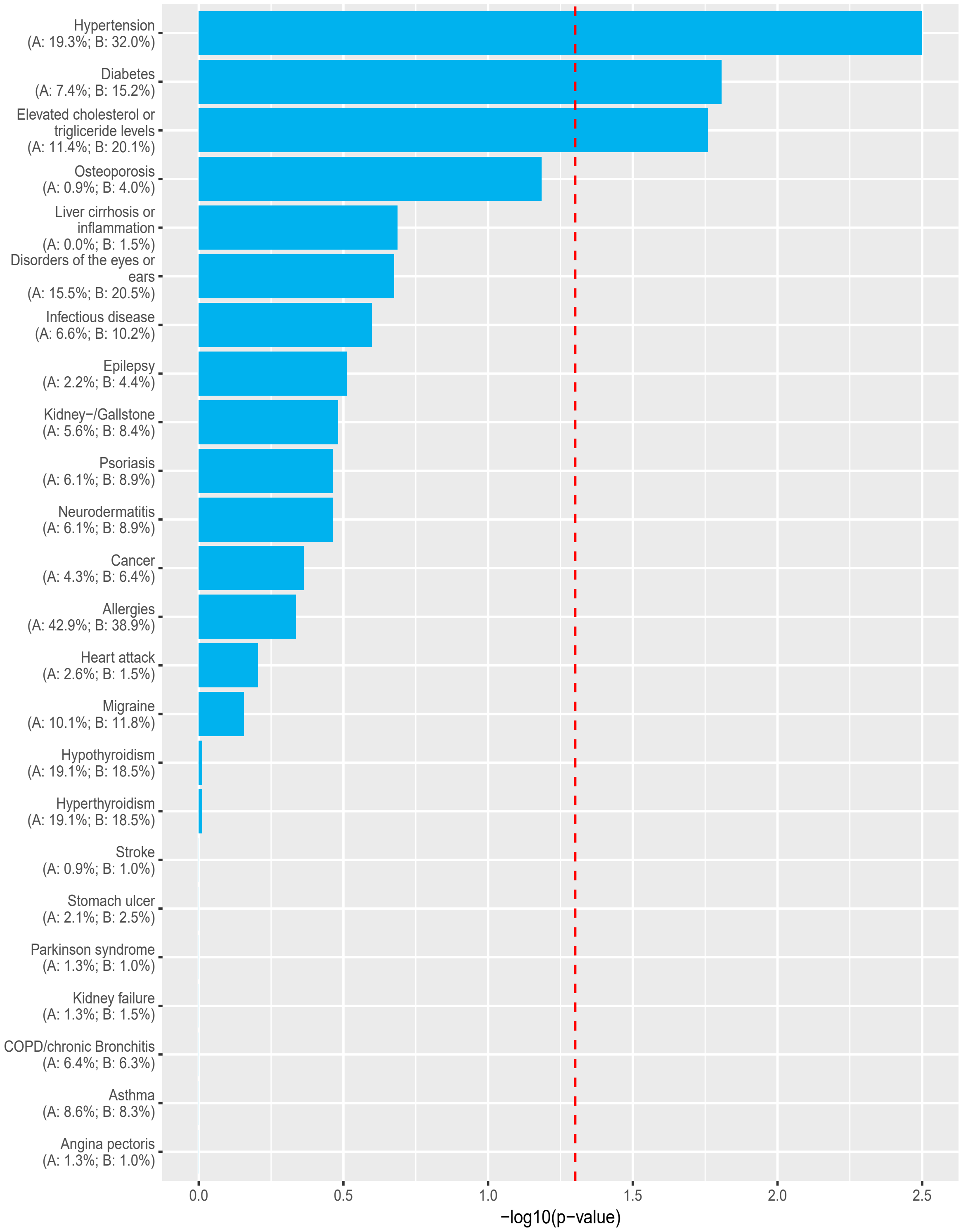

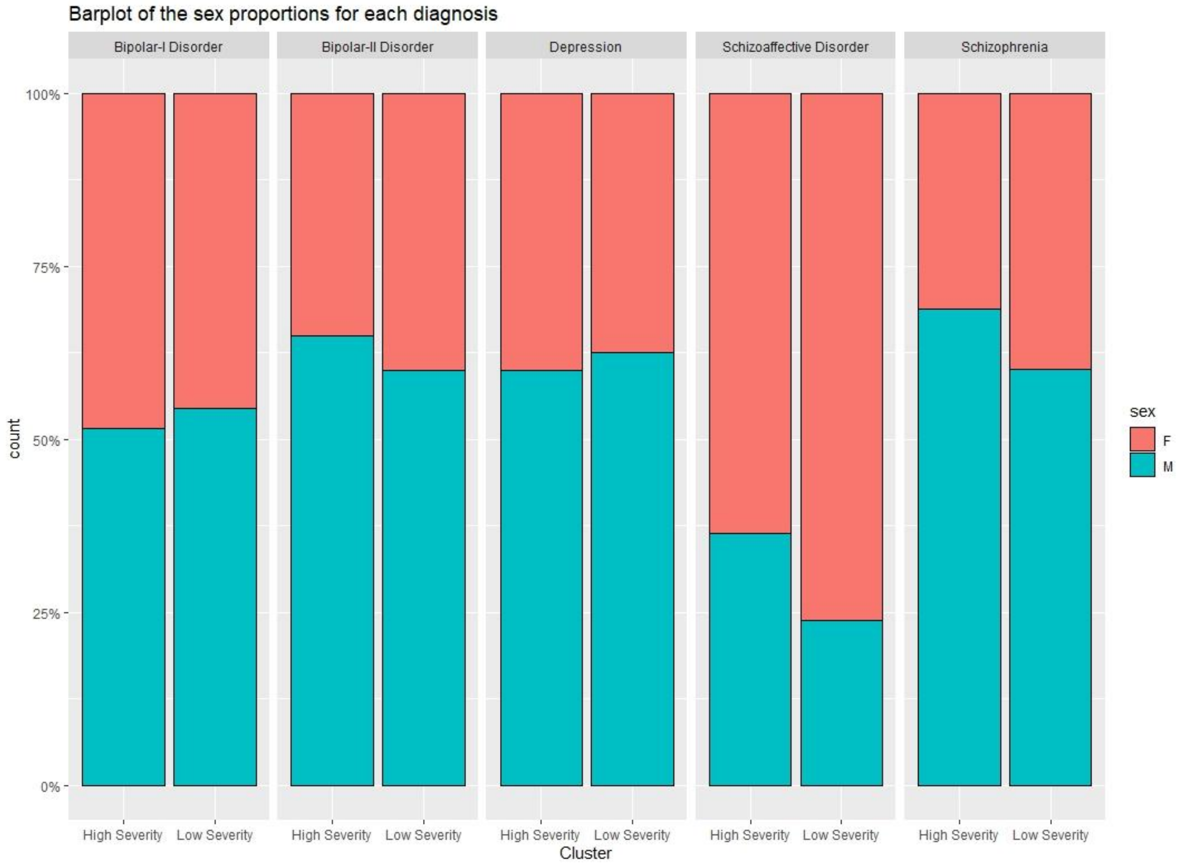

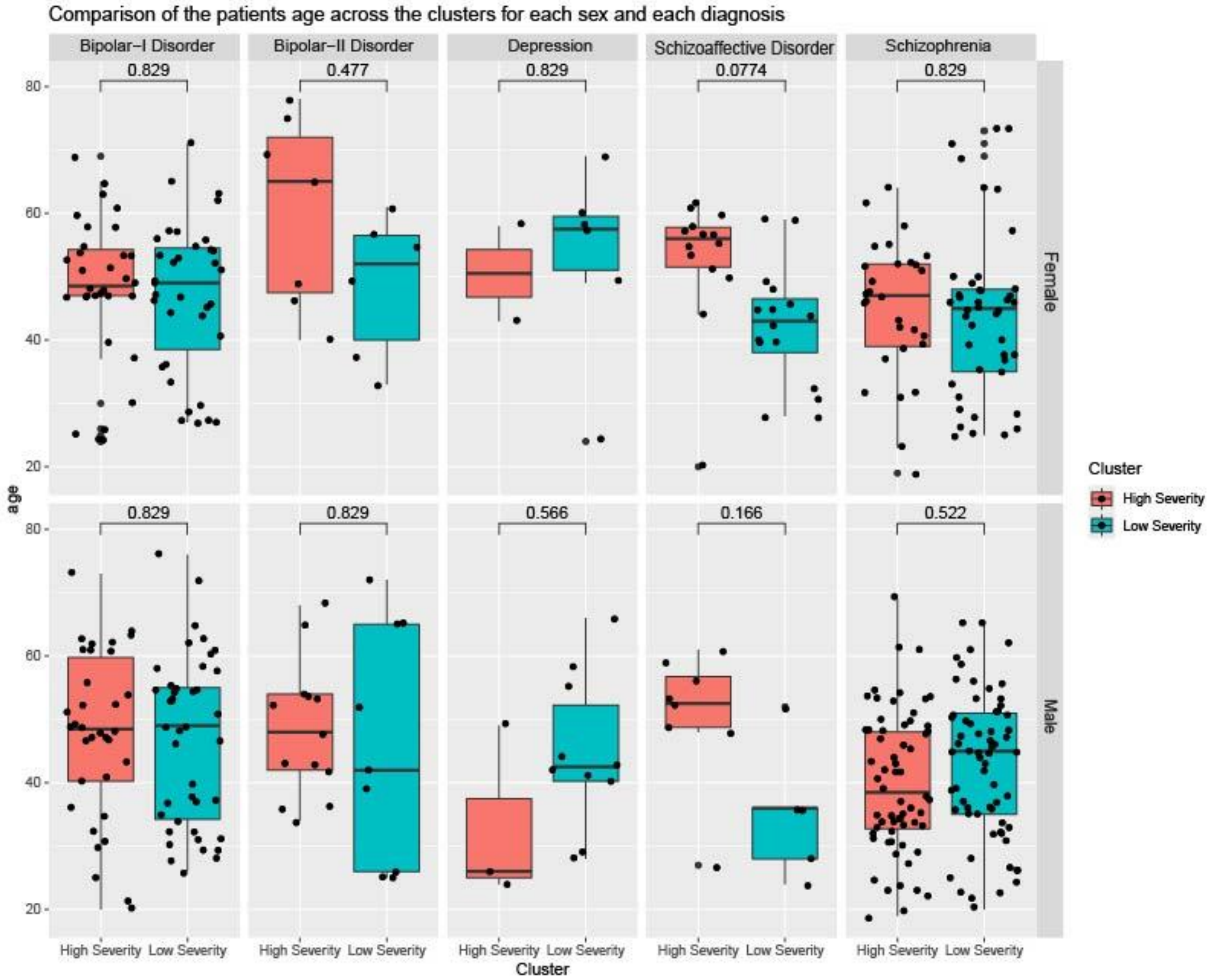

### Supplementary Figure 2

A

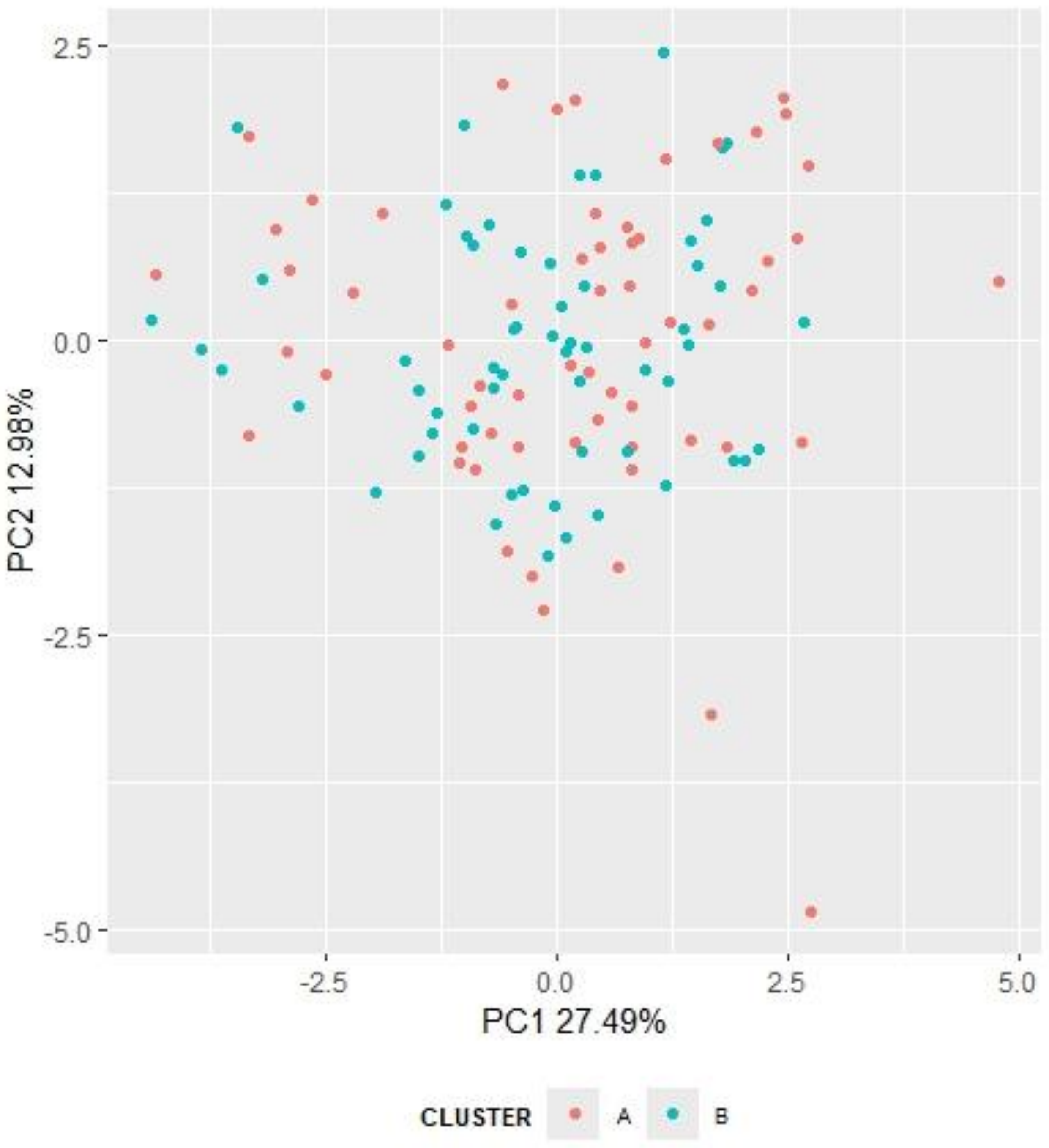

B

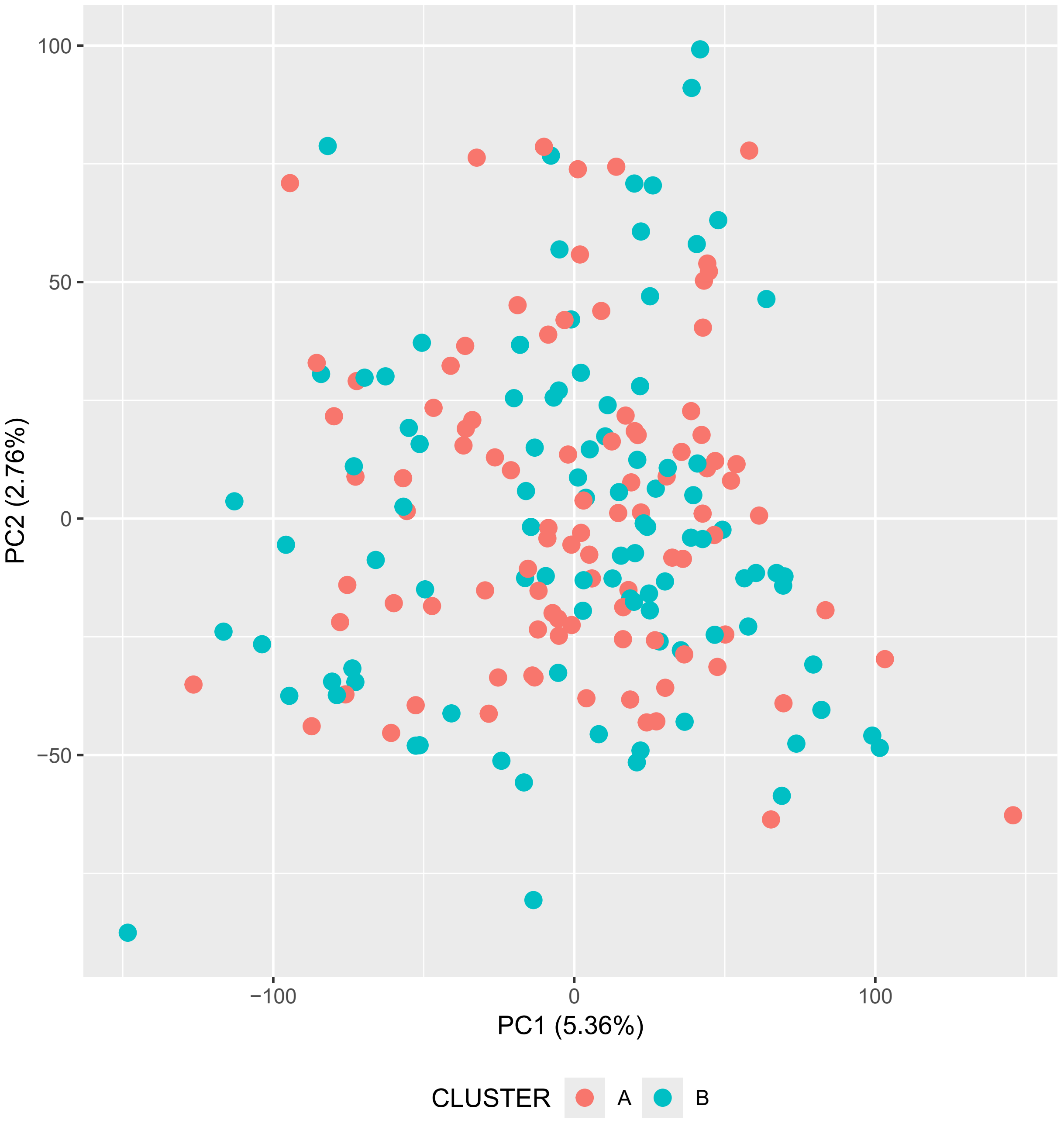

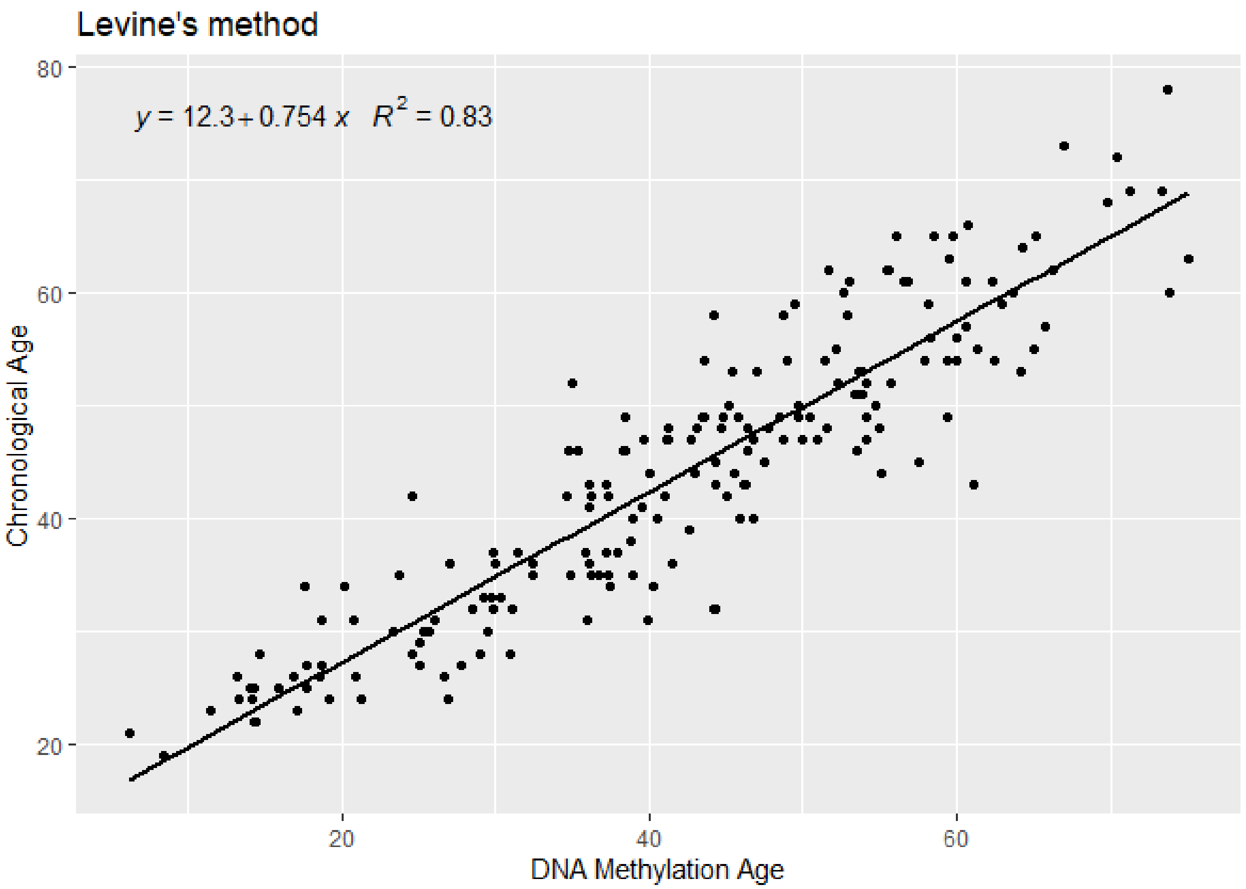

### Supplementary Figure 3

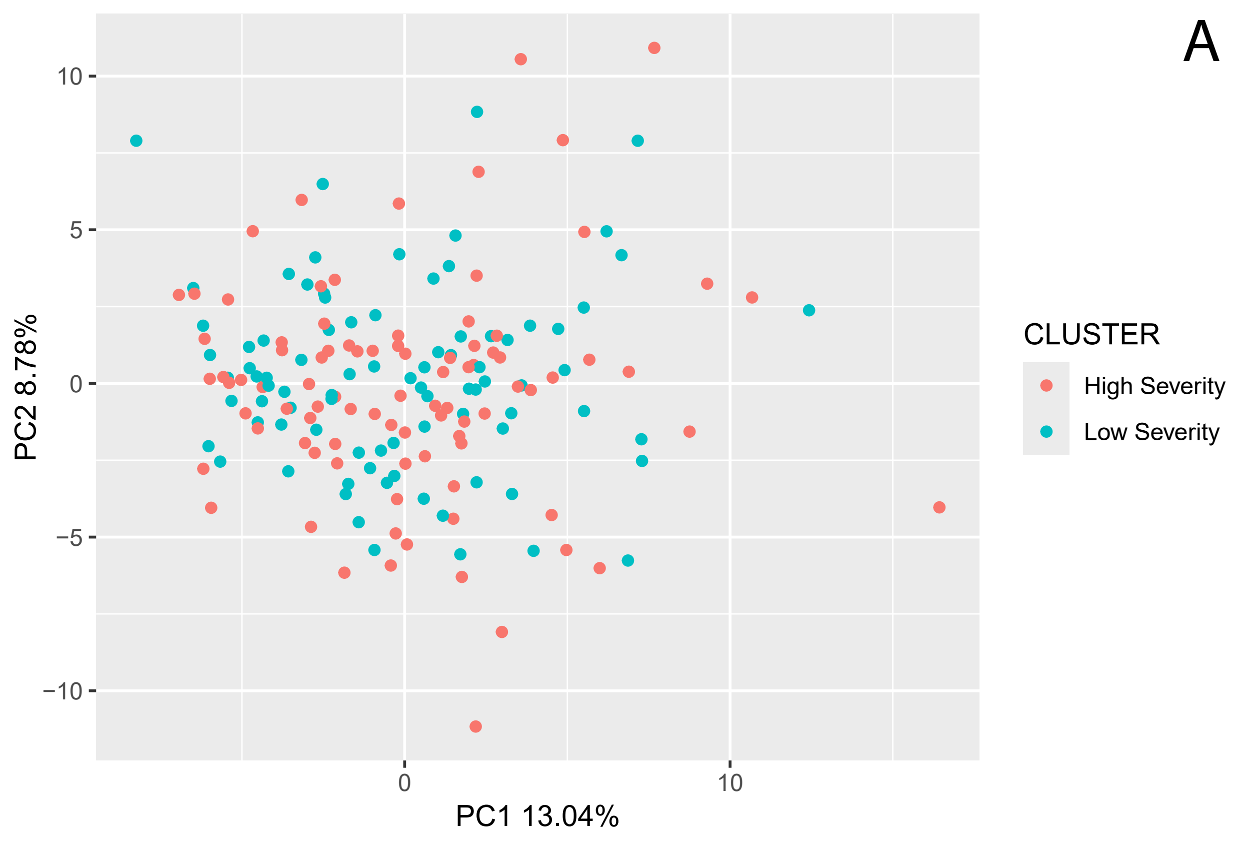

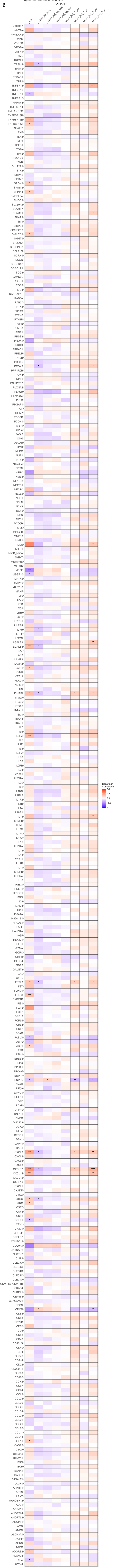

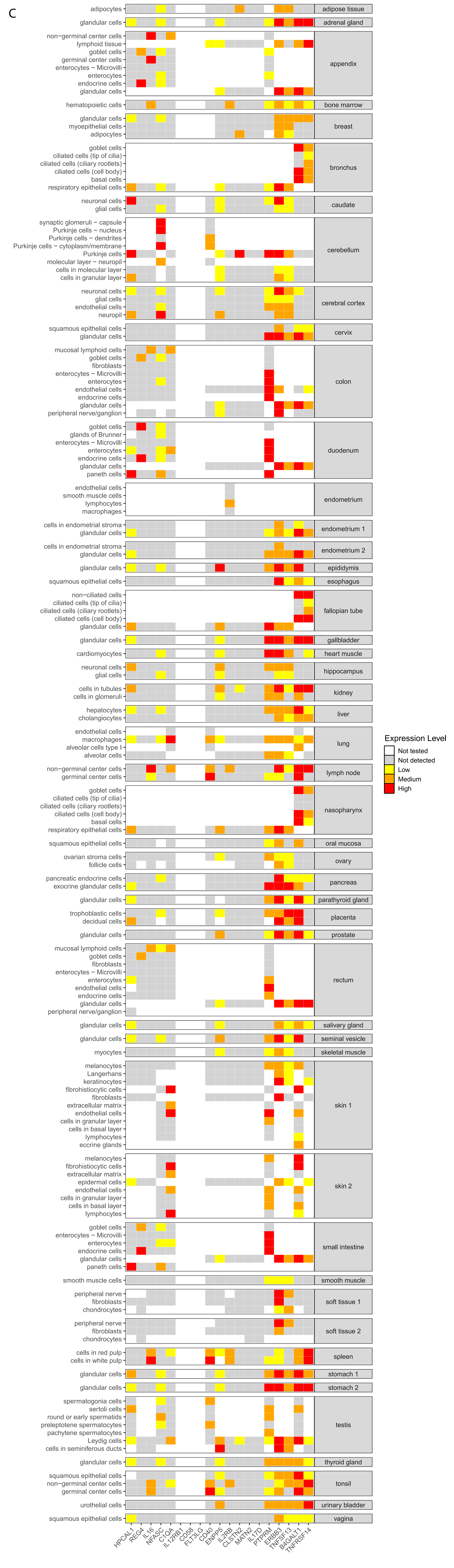

### Supplementary Figure 4

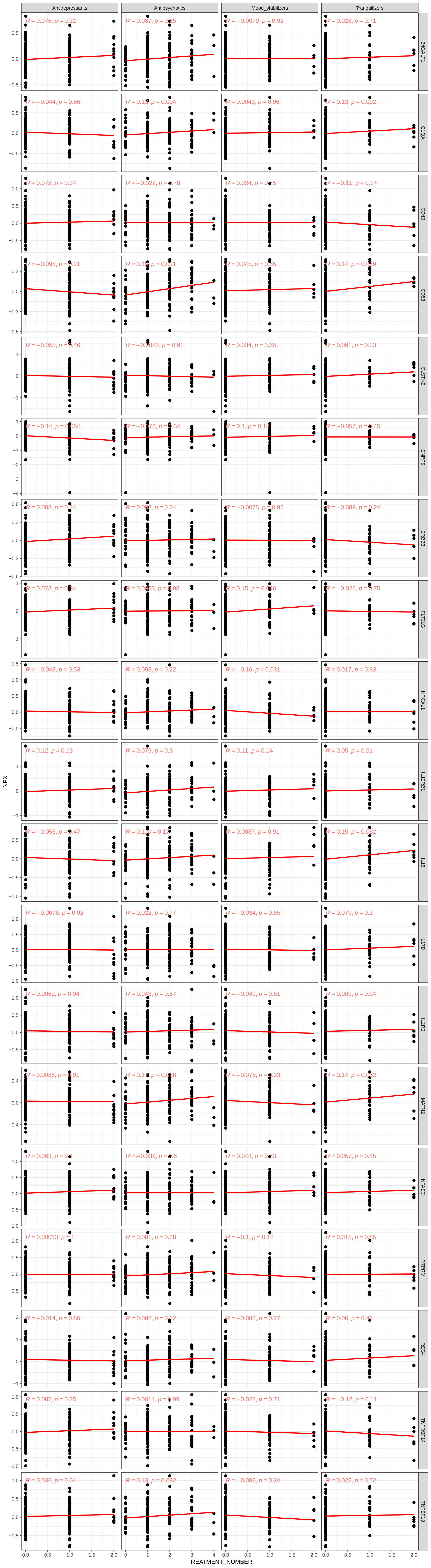
